# CardioMEA: Comprehensive Data Analysis Platform for Studying Cardiac Diseases and Drug Responses

**DOI:** 10.1101/2024.07.28.605490

**Authors:** Jihyun Lee, Eliane Duperrex, Ibrahim El-Battrawy, Alyssa Hohn, Ardan M. Saguner, Firat Duru, Vishalini Emmenegger, Lukas Cyganek, Andreas Hierlemann, Hasan Ulusan

**Author notes:** share senior authorship (Andreas Hierlemann, Hasan Ulusan).

## Abstract

In recent years, high-density microelectrode arrays (HD-MEAs) have emerged as a valuable tool in preclinical research for characterizing the electrophysiology of human induced pluripotent stem cell-derived cardiomyocytes (iPSC-CMs). HD-MEAs enable the capturing of both extracellular and intracellular signals on a large scale, while minimizing potential damage to the cell. However, a gap exists between technological advancements of HD-MEAs and the availability of effective data-analysis platforms. To address this need, we introduce CardioMEA, a comprehensive data-analysis platform designed specifically for HD-MEA data that have been obtained from iPSC-CMs. CardioMEA features scalable data processing pipelines and an interactive web-based dashboard for advanced visualization and analysis. In addition to its core functionalities, CardioMEA incorporates modules designed to discern crucial electrophysiological features between diseased and healthy iPSC-CMs. Notably, CardioMEA has the unique capability to analyze both extracellular and intracellular signals, thereby facilitating customized analyses for specific research tasks. We demonstrate the practical application of CardioMEA by analyzing electrophysiological signals from iPSC-CM cultures exposed to seven antiarrhythmic drugs. CardioMEA holds great potential as an intuitive, user-friendly platform for studying cardiac diseases and assessing drug effects.

## Introduction

As the principal cause of mortality in the Western world, cardiovascular diseases represent a significant healthcare challenge [1]. Moreover, medication intended for non-cardiac conditions can inadvertently induce life-threatening arrhythmias due to off-target effects on the heart [2, 3]. Therefore, preclinical research to investigate drug effects on the heart is of paramount importance. Preclinical testing helps to evaluate the therapeutic potential and to identify any harmful effects of drug candidates targeted at cardiac diseases before commencing human trials.

A combination of human induced pluripotent stem cell (iPSC) and microelectrode array (MEA) technologies has been effectively employed to characterize disease phenotypes and evaluate drug responses *in vitro*. Measurements of either extracellular field potentials [4–7] or intracellular-like signals [8–10] emanating from iPSC-derived cardiomyocytes (CMs) have been conducted. The use of high-density MEAs (HD-MEAs) offers notable advantages in characterizing the electrophysiology of electrogenic cells and conducting drug tests. These advantages include the non-invasiveness of the technique, allowing for long-term recordings with minimal cellular damage, a high signal-to-noise ratio (SNR) [11], the ability to explore characteristics of signal propagation, and the capacity to simultaneously record activities of hundreds of cells at high spatiotemporal resolution [12, 13].

Over the past decade, advancements in micro- and nanotechnology have pushed the boundaries of HD-MEA technology, with an increasing electrode density and number of readout channels generating large data volumes. Despite significant progress in HD-MEA technology, platforms to process and interpret this data remain scarce. Previous studies have offered MATLAB-based software using graphical user interfaces (GUIs) for processing and visualizing MEA data [14, 15]. While GUI applications provide a user-friendly interface and direct interaction with the operating system for efficient computation, they have limitations in the adaptability of the data-processing steps. If users wish to alter processing steps or handle varying file formats, proficiency in GUI programming becomes an essential prerequisite. Additionally, MATLAB is a commercial software, which generates additional costs, when code alterations are necessary. Cardio PyMEA, an open-source application, was recently proposed to address some of these issues [16]. Given its Python foundation, an accessible and extensively utilized programming language, Cardio PyMEA holds the promise to serve a wider user base. It has been designed for ease of GUI element modification, which further increases its appeal.

While there has been considerable progress in the development of analysis platforms for MEA data of CMs, there is still a lot of work to be done to achieve comprehensive characterization of the electrophysiological characteristics of healthy and diseased cells or potential drug responses. Most existing platforms have been designed for working with MEAs that feature a comparably low number of electrodes and offer low spatial resolution, which poses a challenge given the fact that state-of-the-art HD-MEAs feature thousands of readout electrodes and channels [11, 17, 18] and have an electrode pitch as small as 11.47 µm [19]. HD-MEAs generate massive numbers of data points per experiment, so that a statistically relevant sample size can be quickly reached. Moreover, due to the high density and large number of electrodes, they enable reliable signal- conduction-speed estimation of the CM cell assembly [20].

Most of the currently used data-analysis platforms have been designed for processing and analysis of extracellular signals and do not fully leverage the capabilities of HD-MEAs, as recent studies have shown that HD-MEAs are capable of recording not only extracellular but also intracellular-like signals on demand [10, 21, 22]. Compared to intracellular-like measurements, the measurement of merely extracellular activities may not be sufficient to adequately capture drug-induced alterations in cardiac membrane potentials [23]. In most existing platforms, data processing and visualization steps are integrated into a single pipeline. However, this integration leads to inefficiencies in comparative analysis across multiple data files, as each file requires a significant amount of time to process.

For a comprehensive evaluation, an analysis platform should enable the comparison of drug responses at different concentrations or across multiple cell lines with adequate visualization. Furthermore, data processing should be efficient and make optimal use of available computational resources, such as multiple central processing units (CPUs) and memory space, which are often not needed for visualization and comparative analysis. Therefore, the execution of all analysis steps in a single pipeline does not provide optimal performance.

To overcome the aforementioned challenges and limitations, we developed CardioMEA, a comprehensive data analysis platform providing a set of pipelines for the extraction of raw HD-MEA data, feature extraction, data storage, visualization, and advanced analysis (Figure 1). CardioMEA includes data-processing pipelines and a web-based dashboard for data visualization and feature analysis, which have been designed to ensure reproducibility, scalability, and maintainability of all processing tasks. Numerous data files can be processed in parallel using multiple CPUs, which entails a significant reduction in computation time. The resulting processed data are stored in a structured query language (SQL) database, enabling tracking of data history and previous processing steps. Moreover, the CardioMEA Dashboard offers an interactive, web-based platform for data visualization and analysis. This “no-code” application is designed to benefit a broad range of users, facilitating exploratory data analysis, while only a minimal effort is required to understand the underlying code. We demonstrate that - with just a few mouse clicks within the CardioMEA Dashboard - it is possible to analyze CM data of three iPSC lines and their responses to seven antiarrhythmic drugs.

**Figure 1.**
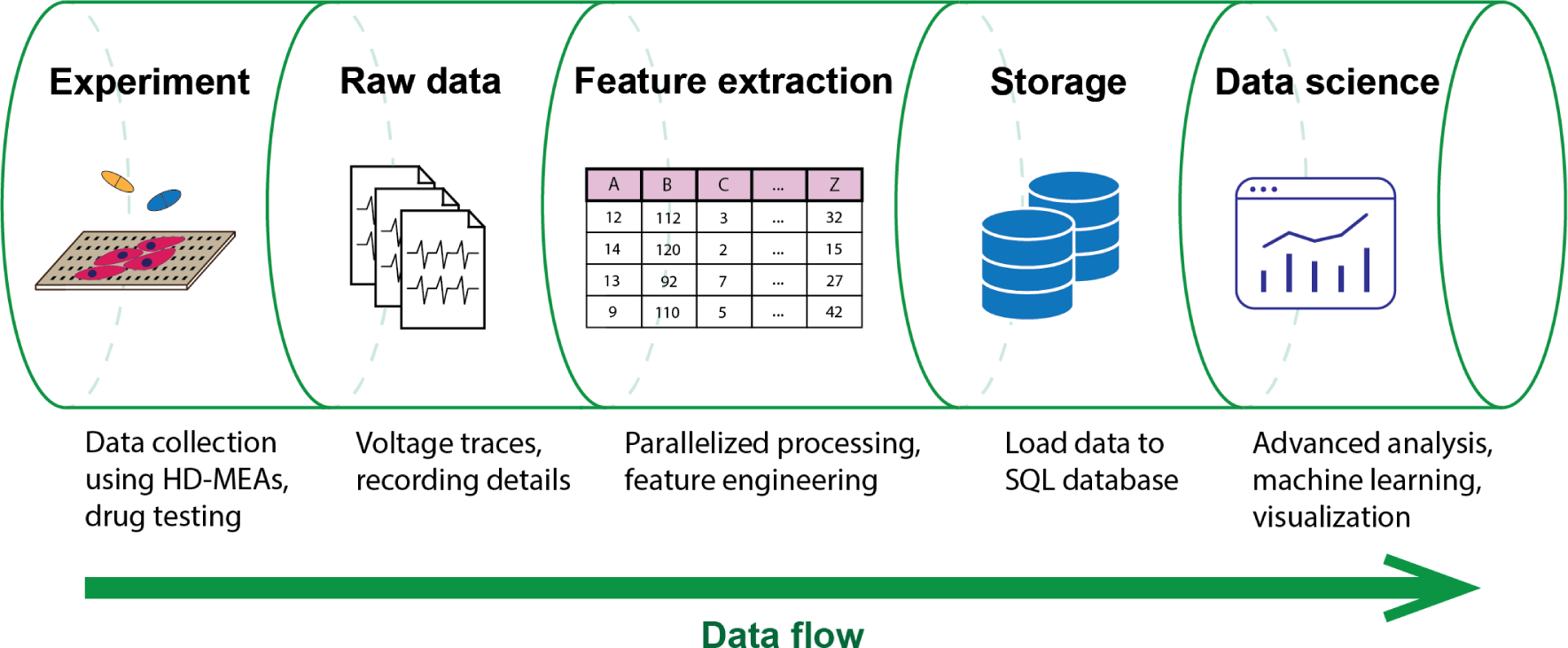
Data flow in the CardioMEA data analysis platform. CardioMEA incorporates every data analysis stage within its structure, ranging from an initial identification of experimental recording files, through raw-data handling and feature extraction, to the subsequent data uploading to an SQL database. Further, it provides a robust system for data visualization and advanced analysis, offering a comprehensive solution for evaluating and interpreting experimental data.

CardioMEA offers unique functionalities for feature analysis, enabling users to assess the predictive power and contribution of each feature in classifying data collected from healthy and diseased cells, as demonstrated in this study. In addition, CardioMEA presents the first open-source platform that can also process and visualize intracellular-like signals of CMs, recorded with HD-MEAs, to obtain detailed insights into membrane potential dynamics [23]. Its open-source configuration and standardized structure may empower a broad spectrum of users in cardiology and the pharmaceutical sector to effortlessly implement and adjust the platform according to their specific requirements.

CardioMEA has been designed for use by scientists with little or no experience in programming, supporting their efforts to investigate the efficacy or potential toxicity of compounds.

As the field of cardiac disease, toxicity research, and individualized precision medicine continues to expand, with ever-increasing data volumes, the need for efficient, scalable, and user-friendly data-analysis tools will grow correspondingly. By efficient streamlining of intrinsic processes and providing an interactive dashboard for data visualization and advanced analysis, CardioMEA will meet these demands and drive advancements in biomedical research and therapeutics.

## Materials and Methods

### Data Science Framework to Build the Data Pipeline

For the development of an open-source data analysis platform, adherence to software engineering best practices is essential to ensure that the code is both readily comprehensible and maintainable. Furthermore, the possibility of reproducing data processing and analysis is a crucial consideration in constructing such a platform. To meet these requirements, we utilized Kedro [24], a Python-based open-source framework renowned for fostering the development of modular and maintainable data science platforms. Kedro comes with built-in wrappers that manage input and output data in diverse formats, including comma-separated values (CSV) and SQL. These features enable efficient data extraction, transformation, and loading processes - critical aspects that make Kedro a suitable tool for developing the data pipelines for this study. By constructing pipelines, which contain nodes chained sequentially, we created a data-flow structure that is both easy to comprehend and reproducible. The principles of clarity and reproducibility were consistently prioritized in our platform’s design and functionality.

### Feature Extraction Algorithms

To extract features from the raw data, we developed two data pipelines to process extracellular and intracellular signal features. An overview of the extracted features and detailed descriptions can be found in Table 1. Among the extracellular signal features, parameters, such as R-wave spike amplitude, R-wave spike width, and field potential duration (FPD) were used as illustrated in Figure 2a.

**Figure 2.**
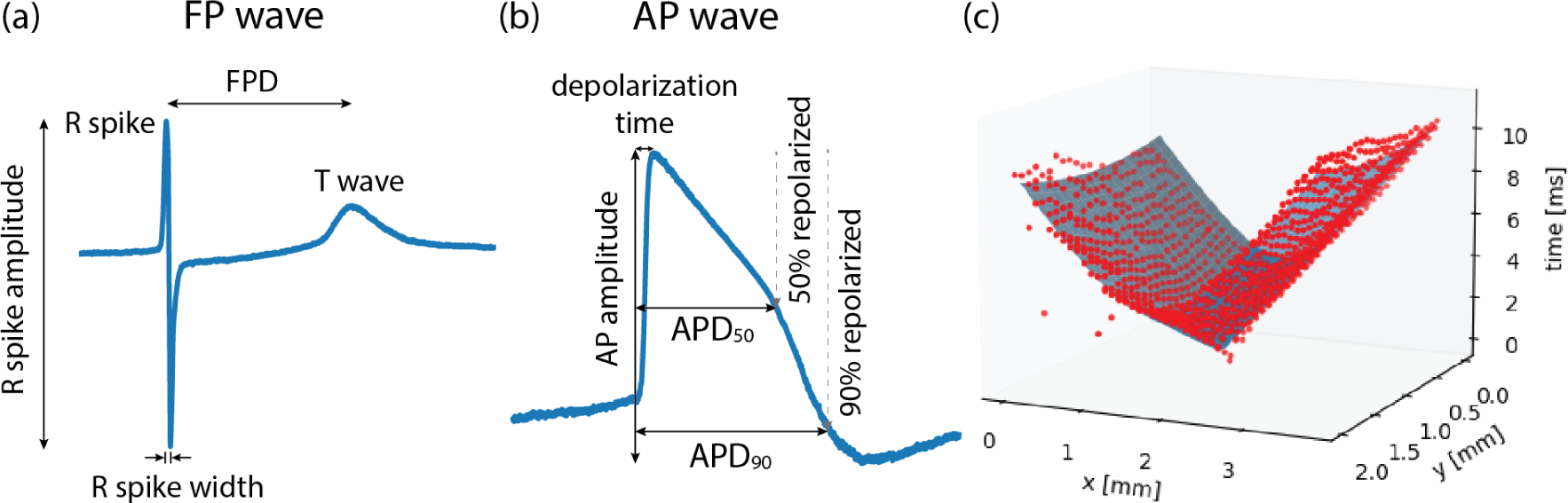
Illustrations of (a) field potential (FP) wave features, (b) action potential (AP) wave features, and (c) conduction speed estimation. FP waves are obtained from extracellular measurements and AP waves from intracellular or intracellular-like measurements. Field potential duration (FPD) is the time difference between the R spike and the T wave. Action potential duration (APD) is the time taken from the onset of depolarization until 50% repolarization (APD50) or 90% repolarization (APD90). (c) To estimate the conduction speed, data points (red color), collected from FP recordings, were fitted to a cone-shaped surface (blue color). Each red dot represents FP data from one electrode.

**Table 1.** Overview of the feature types and names, along with detailed descriptions of the respective computational processes involved in their derivation.

| Feature type | Feature name | Description |
| --- | --- | --- |
| Extracellular signal features | R_amplitude | Difference between the maximum and minimum voltage values of the R spike. |
|  | R_width | Width of the R spike, between the positive and negative peak. |
|  | FPD | Distance between the R spike and the T wave. |
|  | conduction_speed | Mean of local conduction speeds computed at each electrode location. |
|  | rec_duration | Total recording duration in seconds. |
|  | rec_proc_duration | Duration of the processed recording segment. |
|  | n_beats | Number of synchronous beats. |
|  | n_electrodes_sync | Number of electrodes that captured synchronous activity. |
|  | active_area_in_percent | Percentage of electrodes that captured synchronous activity. |
|  | mean_nni | Mean of RR-intervals. |
|  | sdnn | Standard deviation of RR-intervals. |
|  | sdsd | Standard deviation of differences between adjacent RR-intervals. |
|  | rmssd | Square root of the mean of the sum of the squared differences between adjacent RR-intervals. |
|  | median_nni | Median absolute values of successive differences between RR-intervals. |
|  | nni_50 | Number of interval differences of successive RR-intervals larger than 50 ms. |
|  | pnni_50 | Proportion derived by dividing nni_50 by the total number of RR-intervals. |
|  | nni_20 | Number of interval differences of successive RR-intervals larger than 20 ms. |
|  | pnni_20 | Proportion derived by dividing nni_20 by the total number of RR-intervals. |
|  | range_nni | Difference between the maximum and minimum RR-intervals. |
|  | cvsd | rmssd divided by mean_nni. |
|  | cvnni | Ratio of sdnn divided by mean_nni. |
|  | mean_hr | Mean heart rate. |
|  | max_hr | Maximum heart rate. |
|  | min_hr | Minimum heart rate. |
|  | std_hr | Standard deviation of the heart rate. |
| Intracellular signal features | AP_amplitude | Difference between the maximum and the lowest dip of the AP wave. |
|  | depolarization_time | Time from the beginning of the upstroke until the potential reaches the maximum amplitude. |
|  | APD<sub>50</sub> | Time from the beginning of the upstroke until the potential reaches 50% repolarization. |
|  | APD<sub>90</sub> | Time from the beginning of the upstroke until the potential reaches 90% repolarization. |
| Recording information | gain | Gain settings used for the recording. |
|  | cell_line | Name of the cell line. |
|  | compound | Name of the drug, if the recording was made during drug experiments. |
|  | file_path | File path where the recording file was stored. |

The conduction speed was computed following the method described in previous studies [16, 20], with some modifications. In brief, the elapsed time, denoted as T, from the onset of wave propagation, along with the x and y coordinates of the electrode within the MEA, were fitted to a three-dimensional, cone-shaped surface (Figure 2c). This representation can be found in Equation (1).

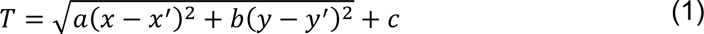

In Equation (1), the coefficients are represented as *a*, *b*, and *c*, while *x*^′^and *y*^′^ are the coordinates of the wave propagation initiation. Following the fitting of the data to the three-dimensional, cone-shaped surface, local conduction velocities at each electrode location were computed as described previously [20]. Subsequently, the magnitudes of local conduction velocities were averaged to derive an estimate of the overall conduction speed.

Among other extracellular signal features, time domain heart-rate-variability (HRV) features were computed using a previously published Python package [25]. The HRV features include metrics, such as mean values and standard deviations of intervals of consecutive R spikes (RR-intervals) [26]. HRV features and other features, such as intracellular signal features (Figure 2b) and recording information, are detailed in Table 1.

### Cardiomyocyte Differentiation and Culture

A human iPSC line UMGi129-A, termed SQT5-line, was derived from a short-QT-syndrome type-5 (SQT5) patient harboring a known variant in the CACNB2 gene. The CACNB2 gene is responsible for encoding L-type Ca^2+^ channels, and variants in this gene have been identified as being linked to short QT syndrome [27, 28]. The SQT5- line’s isogenic control UMGi129-A-1, the SQT5corr-line, was generated after correcting the variant in the CACNB2 gene using ribonucleoprotein-based CRISPR/Cas9. Both the SQT5-line and SQT5corr-line were generated in a prior study [28] and delivered in frozen cryotubes for this research. The iPSC lines were then differentiated into spontaneously beating CMs, following a protocol established in the previous study [28]. Additionally, commercially available CMs, differentiated from healthy donor iPSCs and referred to as iCell Cardiomyocytes, were purchased from Fujifilm Cellular Dynamics International (Madison, Wisconsin, USA). The culturing of these cells was performed according to the manufacturer’s guidelines.

### Cell Plating and Activity Measurement on the HD-MEA

To record the activity of CMs, we used HD-MEAs [11], which were equipped with 26,400 microelectrodes and 1024 readout channels. These HD-MEAs were used to measure both intracellular-like and extracellular signals from CMs [10]. Intracellular-like signals, obtained through the HD-MEAs after electroporation [10], featured action potential (AP) waveforms similar to those recorded by current-clamp patch measurements. In contrast, the signal amplitude was considerably lower [10]. The shape similarity facilitated the extraction of AP wave features, including the action potential duration (APD). Before plating the CMs, the HD-MEAs underwent sterilization through immersion in 70% ethanol, followed by thorough rinsing with deionized water and drying under a laminar flow hood. The electrode array, with a size of approximately 4 × 2 mm^2^, was prepared by coating it with human fibronectin solution (Cat. FC010, Merck KgaA, Darmstadt, Germany) at a concentration of 50 µg/ml. This coating process involved incubation at 37 °C for an hour, providing optimal conditions for cellular adhesion, thereby enhancing the quality of the captured signals.

Following the plating of CMs on the HD-MEAs, the devices were placed in a humidified incubator with 5% CO_2_ to allow for recovery and optimal growth. This environment was maintained for over seven days until spontaneous beating was observed, indicating successful cellular adaptation and functioning. The cell culture medium was refreshed three times a week to maintain cell health and vitality during measurements.

The activity measurements on the HD-MEAs were consistently conducted within a humidified incubator set at 37 °C with 5% CO_2_. The controlled environment ensured reproducibility and reliability of our cellular activity measurements. For the extracellular measurements, cellular activity was screened over the entire electrode array area using the MaxLive Software (version 19.2.27, MaxWell Biosystems AG, Zurich, Switzerland). Thereafter, 1020 electrodes featuring the largest signal amplitudes were selected. A similar process was applied to the intracellular measurements, with cellular activity screened and 200 electrodes with the largest signal amplitudes selected. We then performed electroporation on these 200 electrodes to measure intracellular-like signals according to the procedure that has been previously described [10].

### Drug Testing Protocols

Dimethyl sulfoxide (DMSO, Cat. D4540), quinidine (Cat. 22600), nifedipine (N7634), flecainide (Cat. F0120000), amiodarone (Cat. A8423), sotalol (Cat. S0278), and disopyramide (Cat. D2920000) were purchased from Merck KGaA (Darmstadt, Germany). Ivabradine (Cat. HY-B0162A) and ranolazine (Cat. HY-B0280) were purchased from Lucerna-Chem AG (Lucerne, Switzerland). All drug solutions, except ivabradine and sotalol, were prepared in a two-step procedure. The initial dissolution of the compounds was carried out in DMSO, followed by a subsequent dilution in the culture medium. This procedure was specifically designed to ensure that the DMSO concentration remained below 0.1% during drug testing. Ivabradine and sotalol were dissolved directly in the culture medium. For the SQT5-line, drug concentrations in the HD-MEAs were gradually increased using an additive approach to investigate the effects of escalating doses.

We explored the influence of quinidine, flecainide, disopyramide, ivabradine, amiodarone, sotalol, and ranolazine on CMs differentiated from the SQT5-line iPSCs. Following three to four baseline measurements, taken at 30 minute intervals, the drug concentration was sequentially increased as specified in Table 2, with each increment separated by 45-minute intervals, unless otherwise specified in respective figures. Extracellular signals were recorded at each concentration level and stored in standard HDF5 format using the MaxLive Software.

**Table 2.** List of drugs and their concentrations used in the drug testing experiments.

| Recording type | Cell line | Drug | Concentration(s) ($\mu\text{M}$) |
| --- | --- | --- | --- |
| Extracellular recording | SQT5-line | Quinidine | 0.1, 0.3, 1, 3, 10 |
|  |  | Flecainide | 0.03, 0.1, 0.3, 1, 3 |
|  |  | Disopyramide | 3, 13, 43 |
|  |  | Ivabradine | 0.3, 1, 3 |
|  |  | Amiodarone | 0.03, 0.1, 0.3, 1, 3 |
|  |  | Sotalol | 3, 10, 30, 100, 300 |
|  |  | Ranolazine | 0.3, 1, 3 |
| Intracellular-like recording | iCell Cardiomyocytes | Nifedipine | 0.1 |
|  |  | Quinidine | 1 |
|  |  | Sotalol | 30 |

We also investigated the impact of nifedipine, quinidine, and sotalol on the intracellular signals of iCell Cardiomyocytes using electroporation. The applied drug concentrations are specified in Table 2. After establishing three baseline measurements at one-hour intervals, a pre-diluted drug solution was administered to the HD-MEA 30 min after the latest baseline measurement to reach the respective concentration. Intracellular-like signals were subsequently recorded 30 min after the addition of drugs.

## Results and Discussion

### Modular and Structured Data Analysis Pipeline

We developed a data analysis platform, named “CardioMEA”, using the Kedro framework to process and analyze data collected from CMs on HD-MEAs. Our approach ensured modularized data processing so that each step is clear and understandable. Our code’s modular structure treats functions as building blocks that are assembled in a specific sequence to construct the respective pipeline. All pipelines developed within this study are listed in a file named ‘pipeline_registry.py’. This comprehensive record is intended to provide easy access to each pipeline, simplifying navigation and usage throughout the data analysis.

Details pertaining to input and output data, such as the paths to their respective storage locations and configurations for data loading and saving, are cataloged in a dedicated file, called ‘catalog.yml’. This arrangement is particularly advantageous, as it simplifies the management of inputs and outputs without the need to navigate through the entire code.

To use CardioMEA, the initial requirement is the installation of Conda [29], one of the most prevalent package and environment managers amongst Python users. CardioMEA is publicly accessible on GitHub via the following link: https://github.com/leejheth/CardioMEA. Setting up the working environment is straightforward, requiring only the execution of the ‘make setup’ command in the terminal. This command will automatically generate a new Conda environment named ‘cardio-env’ and install all necessary dependencies within it. This user-friendly setup ensures easy access to our sophisticated data-processing tools for a broad community of researchers.

### Feature Extraction from Multiple Data Files in Parallel

Experiments often result in a multitude of recording files requiring specific processing. Undertaking this task for each file can be cumbersome and time-consuming. Therefore, we designed CardioMEA with the capability to handle and process multiple data files concurrently, using multiple CPUs. Users can predetermine the number of CPUs to scale the process, depending on the resource availability of their workstations or high- performance computing clusters.

Initially, the user needs to supply a CSV file containing a list of directories housing the recording files, along with corresponding cell line names, compounds (drugs), and additional notes as applicable. These notes may include any relevant experiment details, such as compound concentrations or experiment IDs. Subsequently, by executing the ‘list_rec_files’ pipeline, a complete list of recording files, stored in the specified directories, can be identified and listed in a newly created CSV file. The recording files, listed in this CSV file, are then processed in batches, with the batch size being equivalent to the number of CPUs predetermined by the user. This approach allows for efficient parallel processing and significantly reduces the time required to process large data volumes.

The subsequent stage involves feature extraction from the recording files. The feature extraction pipelines for extracellular and intracellular data are illustrated in Figure 3. The information for the extracellular data is extracted from the recording file, followed by identifying the spike time points. Then, FP wave features, conduction speed, and HRV features are computed. Similarly, intracellular data extraction occurs from the recording file, which is then followed by the calculation of AP wave features. All extracted feature values, coupled with a timestamp indicating when the processing was completed, are uploaded to a PostgreSQL database. Two distinct SQL tables are utilized, each for extracellular and intracellular data. Each processing result is inserted as a row in the SQL table accompanied by a timestamp, allowing for comprehensive data history preservation within the database. This feature proves particularly beneficial in scenarios where processing steps need to be modified or altered. It enables the tracking of previously processed data, thereby ensuring traceability of all data transformations.

**Figure 3.**
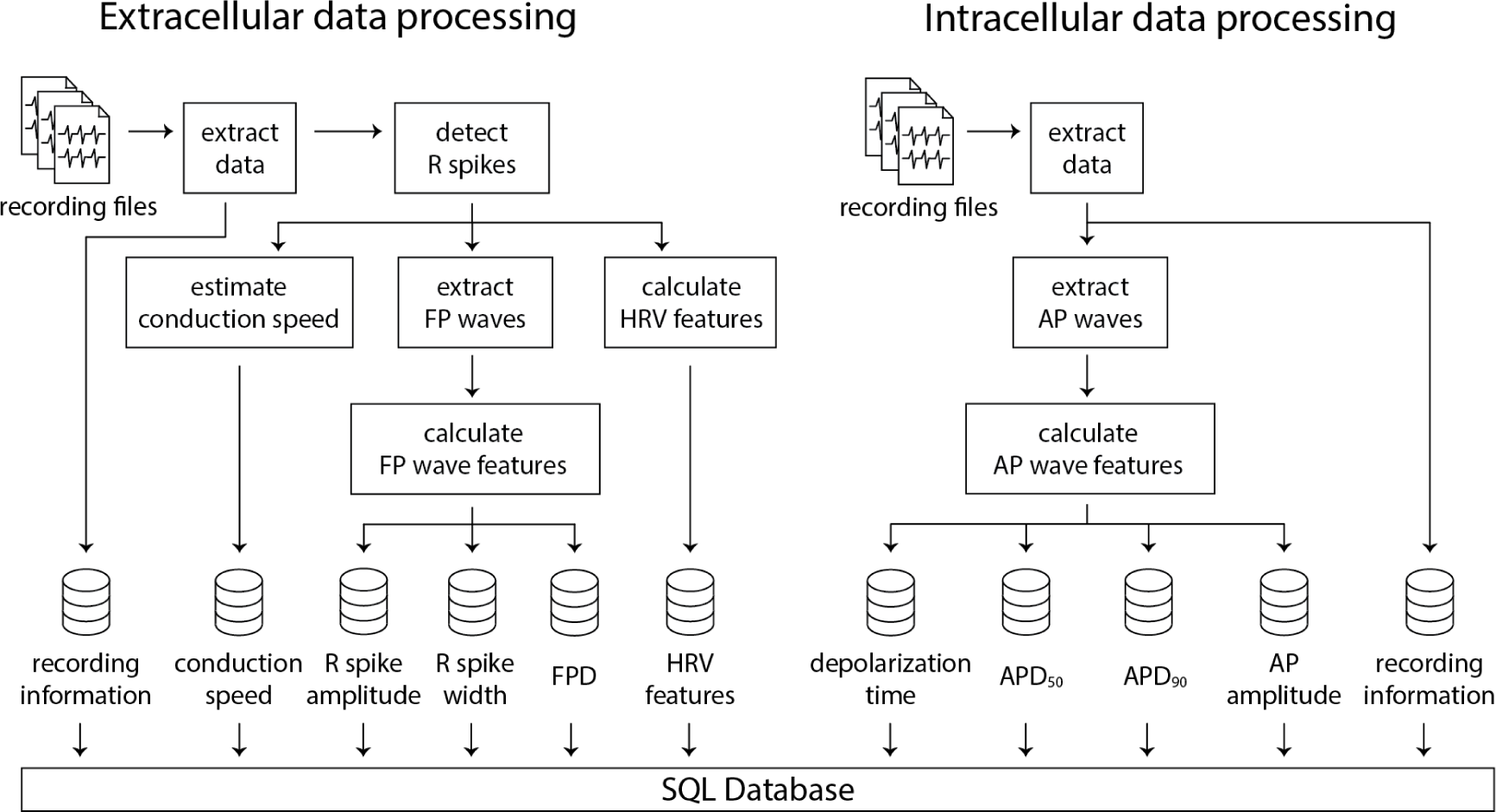
Illustration of the data processing pipelines for extracellular and intracellular data obtained using HD-MEAs. Numerous recording files can be processed in parallel using multiple CPUs, which significantly decreases computation time and enhances efficiency.

### Feature Correlations between Different Types of HD-MEA Data and Whole-Cell Current-Clamp Patch Measurement Data

As CardioMEA cannot only process extracellular signals but also intracellular-like signals from iPSC-derived CMs, researchers can study the relationship between these two types of electrical signals (extracellular and intracellular-like) recorded by the very same electrodes and originating from the same cells. Such an analysis offers a unique perspective on the relationship between extracellular and intracellular or intracellular-like signal and waveform features. We conducted a correlation analysis between features derived from the two different data types. Utilizing data from 29 CM cultures (iCell Cardiomyocytes), we analyzed signals obtained from a total of 3,987 electrodes, which captured both extracellular and intracellular-like signals. Figure 4 presents the correlations between features extracted from the two signal types.

**Figure 4.**
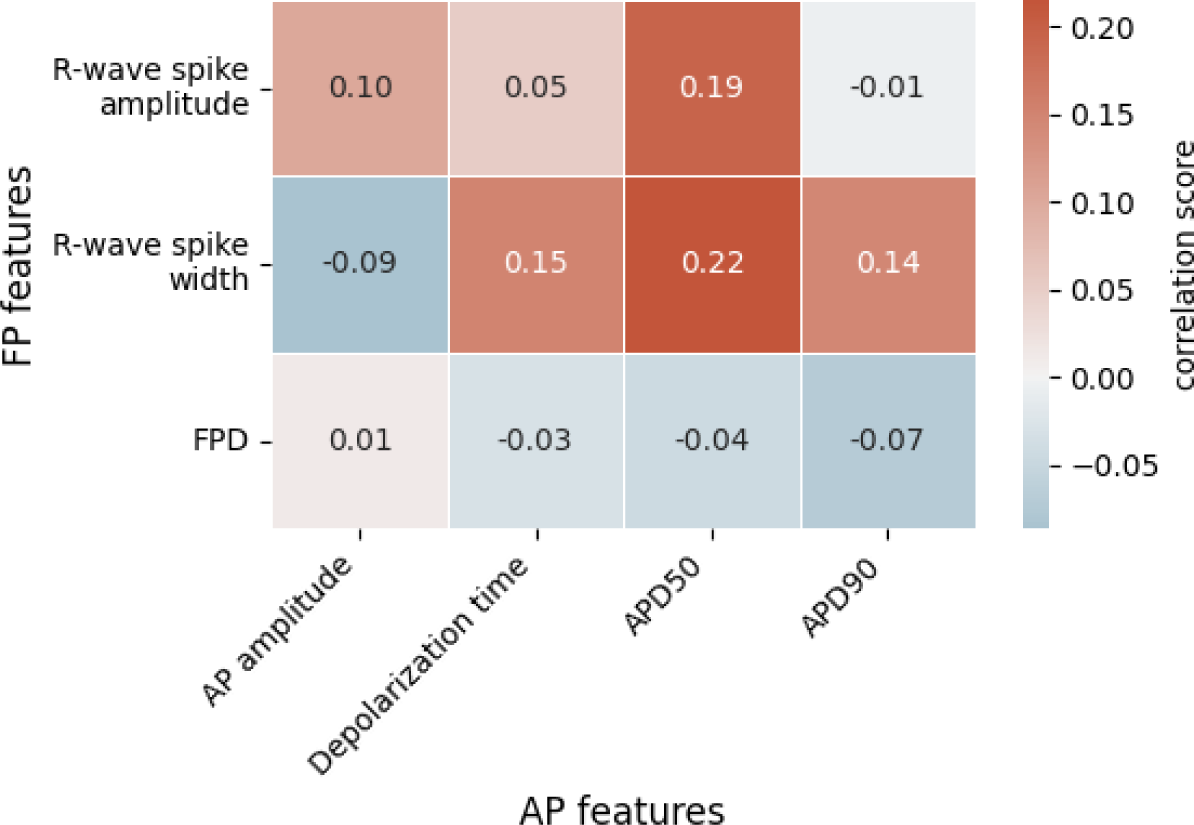
Correlations between extracellular field potential (FP) and intracellular-like action potential (AP) features. Each value represents the Spearman rank correlation coefficient. FPD, field potential duration; APD50, action potential duration at 50% repolarization; APD90, action potential duration at 90% repolarization.

In Figure 4, we see a positive correlation between R-wave spike width and depolarization time, APD_50_, and APD_90_. The R-wave spike appears in the depolarization phase of an AP, which supports a correlation between the R-wave spike width and the AP’s depolarization time. Both APD_50_ and APD_90_ are measured from the start of depolarization and are, therefore, connected to the R-wave spike width. Among the FP features shown in Figure 4, the R-wave spike amplitude is the only feature expressed in terms of voltage; in contrast, R-wave spike width and FPD are time-based measures. Regarding the R-wave spike amplitude, we noticed that R-wave spikes were clipped in some channels due to their high signal magnitude. While the gain may be reduced during recordings to address this clipping issue, it entails the risk of diminishing the T-wave amplitude, which is already small. Consequently, the R-wave spike amplitude is not an ideal feature for correlation analysis.

Interestingly, FPD did not strongly correlate with either APD_50_ or APD_90_, even though we anticipated a relationship with these two parameters. In fact, FPD, in our correlation analysis, showed minimal correlation with all AP-related features. This could be attributed to the noise levels in the FPD data that occasionally obscured the detection of the T-wave. We noted that the T-wave and its exact timing often could not be precisely determined, especially for extracellular signals of iCell Cardiomyocytes featuring very small T-wave amplitudes. Therefore, extracting FPD values and calculating their correlations to other features is challenging.

Next, we derived intracellular-like features from HD-MEA data and whole-cell current clamp patch data to examine the correlation between the respective feature values. For this analysis, we used recordings by patch clamp and intracellular-like recordings by the HD-MEA, which were obtained simultaneously from the same cells. The data were published [10] in a previous study (cell A, Figure S 1a; cell B, Figure S 1b). Figure 5 shows the correlation between features extracted from 60 AP waveforms over a span of 124 sec (cell A) and from 23 AP waveforms over a span of 36 sec (cell B) that have been captured through simultaneous patch clamp and HD-MEA recordings.

**Figure 5.**
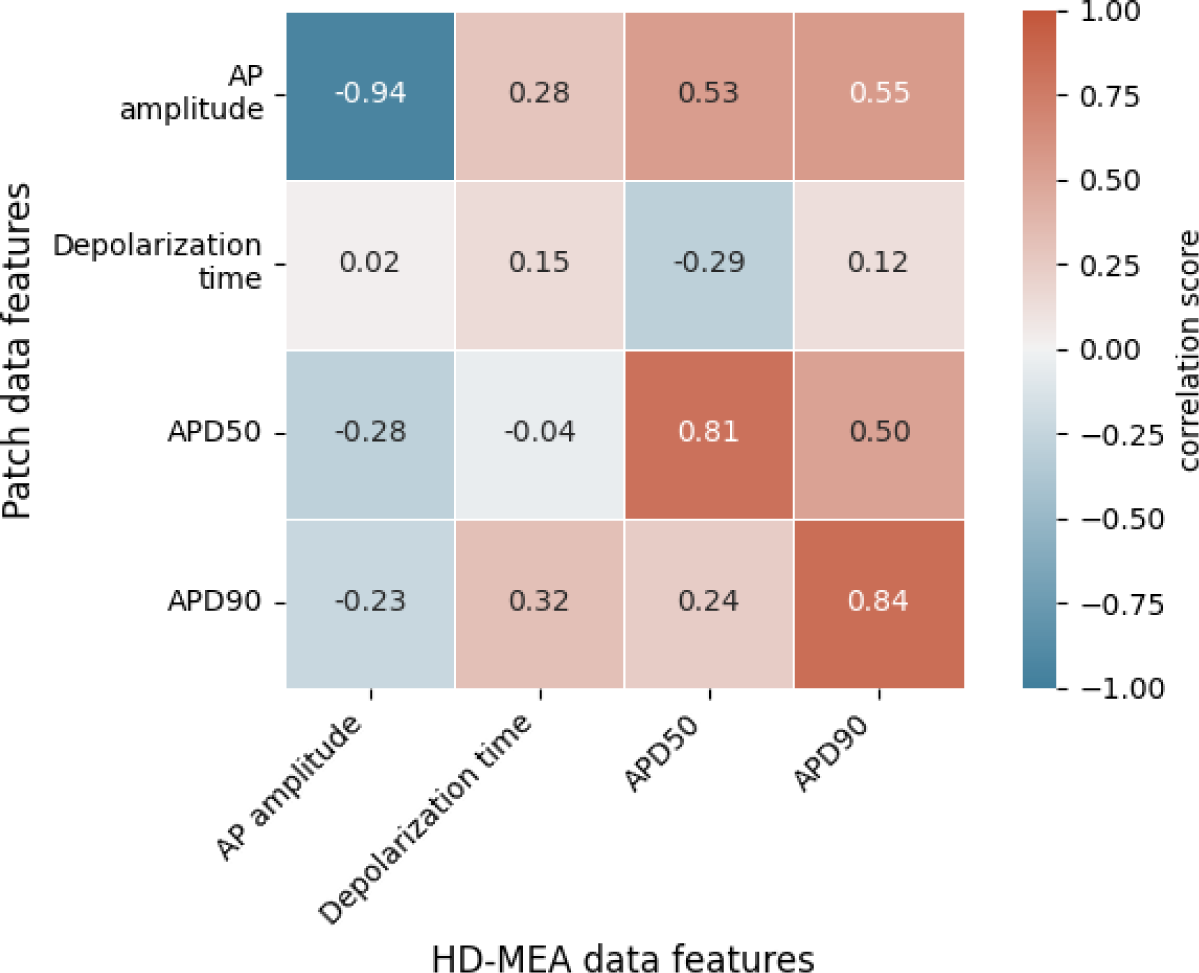
Correlations between intracellular features obtained with patch clamp measurements in current-clamp mode and features of intracellular-like data obtained by HD-MEAs in simultaneous measurements from the same cells. A total of 83 AP waveforms collected from 2 cells (cell A, cell B) were used to compute the correlation scores. Each value represents the Spearman rank correlation coefficient. APD50, action potential duration at 50% repolarization; APD90, action potential duration at 90% repolarization.

Figure 5 shows that AP amplitudes of the patch clamp and the HD-MEA data are negatively correlated. This negative correlation is, however, a consequence of the fact that both signals are simultaneously measured from the same cell(s). As discussed in our previous study [10], ion and current leakage through nanopores, which are generated transiently by the electroporation, reduce the patch clamp signal. The poration-induced leakage decreases over time as the cell membrane reseals. The resealing of the cell membrane increases the AP amplitude of the patch clamp recording, while the AP amplitude in the HD-MEA measurement is concurrently decreasing (Figure S 1). APD_50_ and APD_90_ exhibited strong positive correlations between patch clamp and HD-MEA measurements, indicating that the two recording methods capture consistent APD patterns over time and while the cell membrane reseals. Depolarization times, on the other hand, were only weakly correlated between the two recording methods. This finding may be due to differences in waveform shapes, as shown in Figure S 1 and Figure 6. In particular, the AP waveforms, captured by HD-MEA measurements, exhibit a gradual increase of the voltage before the rapid upstroke (before Phase 0) which is much less pronounced in the AP waveform captured by the patch clamp measurement. As the data are collected simultaneously from the same cells, this marked difference may be attributed to the difference in the two recording settings, most prominently the electrode configuration (penetrating patch pipette and outside Pt-black coated planar electrode).

**Figure 6.**
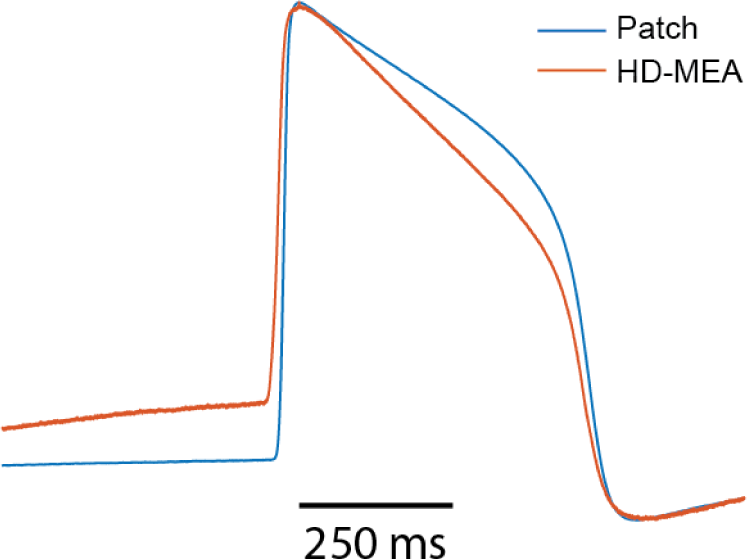
Overlay of amplitude-normalized AP waveforms, averaged across 60 consecutively recorded waveforms (cell A) and obtained from patch-clamp and HD-MEA measurements.

Interestingly, the AP waveforms captured by the HD-MEA closely resemble those from previous studies [8, 30, 31] that reported electroporation-mediated intracellular-like recordings with planar electrodes on MEAs. Conversely, when sharp 3D nanostructures were employed for intracellular-like measurements, AP waveforms obtained through MEAs more closely resembled those of patch clamp measurements [32, 33]. These observations suggest that the difference in AP waveform shapes between patch-clamp and HD-MEA recordings may be attributed to differences in electrode shape and arrangement, which likely influence the correlation between depolarization time values (Figure 5) obtained from both methods.

As becoming evident from this section, CardioMEA’s unique features and unprecedented capability of processing and analyzing both extracellular and intracellular-like signals, captured by HD-MEAs, offer invaluable insights. These insights enhance our understanding of cardiac electrophysiology and facilitate correlation analyses between different data types.

### Interactive Dashboard for Exploratory Data Analysis and Visualization

After data processing, the availability of an interactive tool that enables scientists to visualize the processed data and conduct exploratory data analysis is essential for data-driven analysis. Therefore, we have incorporated a web-based interactive dashboard within CardioMEA, termed “CardioMEA Dashboard”. This user-friendly interface facilitates the visualization of data in the SQL database, further investigations of features, and downloading the resulting figures. Furthermore, the web-based nature of the CardioMEA Dashboard makes it a very accessible tool. Unlike traditional GUI-based software, the CardioMEA Dashboard does not require specific installation, thereby offering enhanced compatibility with a broad range of user operating systems. This feature ensures universal applicability and ease of use, rendering the CardioMEA Dashboard a reliable and convenient tool for biomedical researchers working in diverse computational environments.

The dashboard’s data panel, illustrated in Figure 7, displays all cell lines and compounds found in the existing SQL database, either from the extracellular or intracellular data table. Upon choosing specific cell lines and compounds, data corresponding to the selected criteria will be presented in the ‘List of processed files’ table. Users can then select multiple files of interest, which will be displayed at the bottom of the data panel. This arrangement empowers users to navigate the SQL database and interactively select data for analysis without the need of proficiency in SQL.

**Figure 7.**
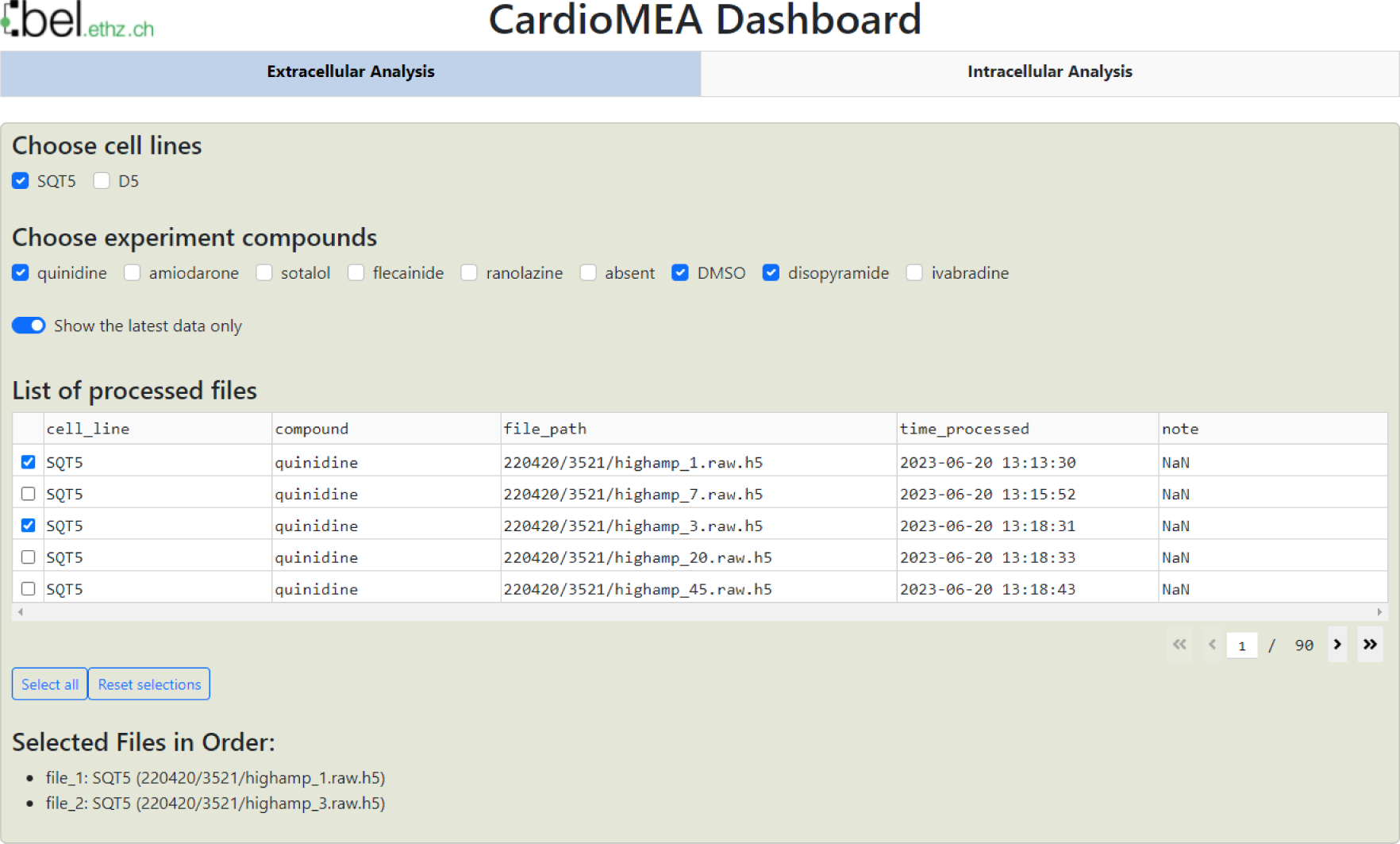
CardioMEA Dashboard data panel. The cell lines stored in the SQL database appear at the top row of the data panel, while the compounds, used with each selected cell line, are shown in the subsequent row. Users have the option to display either all historical processed data or limit the display to the most recent data. Based on these settings, processed files are listed in the table, which allows for selecting specific data for visualization and further analysis.

Located beneath the data panel is a visualization and analysis panel divided into three tabs (see also Figure S 10). The first tab, titled ‘Data distribution’, contains a set of figures visualizing the data. As an example for extracellular data analysis, Figure 8 represents the evolution of the R-wave spike amplitude, the R-wave spike width, FPD, and the conduction speed during an increment of quinidine concentration dosed to CMs, differentiated from SQT5-line iPSCs, after four baseline measurements. Quinidine is a Class Ia antiarrhythmic drug that is known to prolong cardiac repolarization [34, 35]. In clinical investigations involving short-QT-syndrome patients, quinidine effectively prolonged the QT interval [36, 37]. Other studies have reported that quinidine prolonged APDs of CMs derived from short-QT-syndrome type-1 patients [38, 39]. When quinidine was applied to CMs derived from an SQT5 patient, we observed an increase in FPD with increasing quinidine concentration (Figure 8), which is in agreement with findings from a previous study [28], suggesting that quinidine may also effectively prolong the QT interval of SQT5 patients. Additionally, a decrease in signal-conduction speed was observed as the quinidine concentration increased. This reduction in conduction velocity can be attributed to quinidine’s blocking effect on Na^+^ channels. By limiting the influx of Na^+^ ions into the cell, quinidine may indirectly slow down the diffusion of Na^+^ ions to adjacent cells that are interconnected through gap junctions.

**Figure 8.**
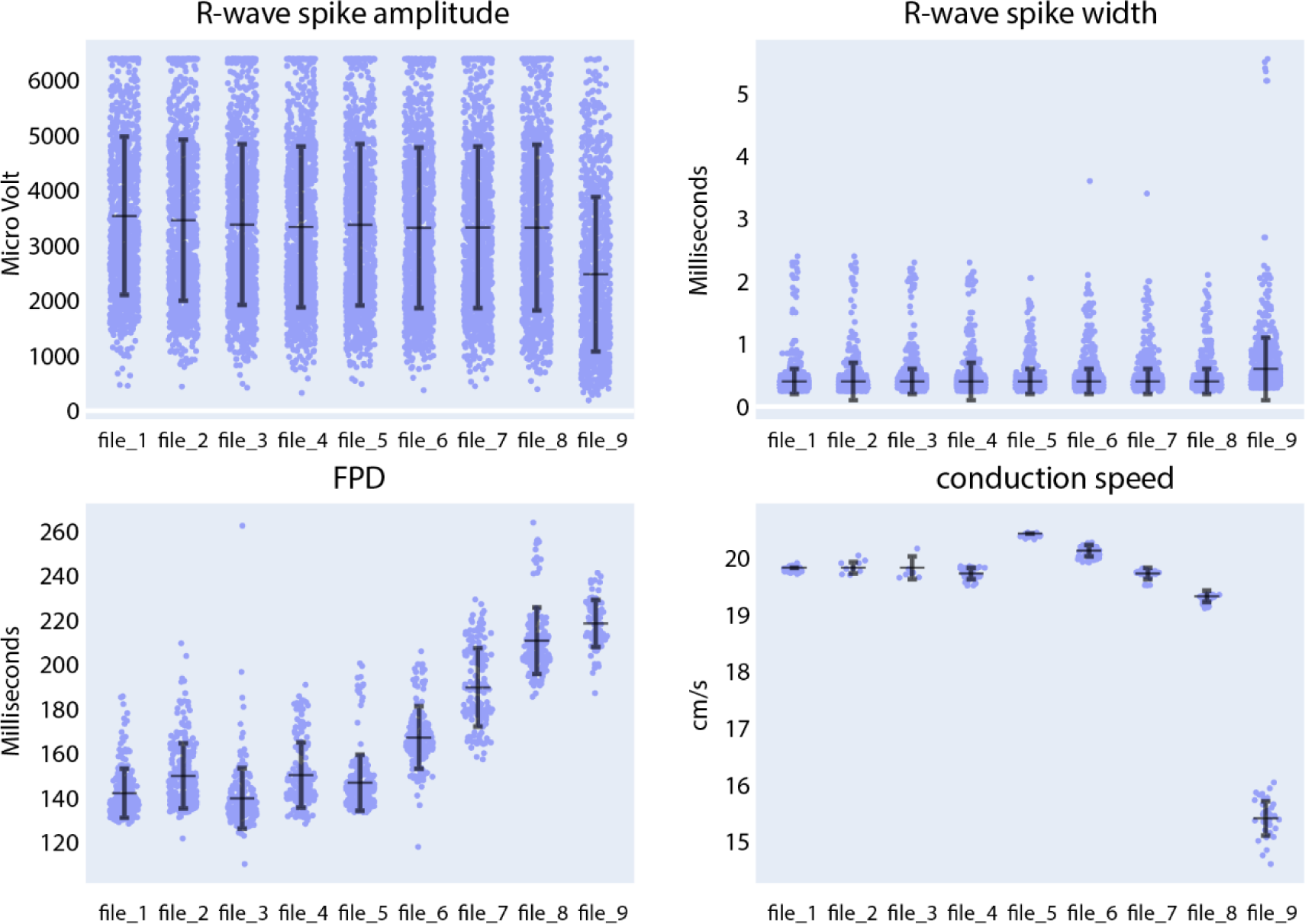
Compound analysis of signals of CMs derived from SQT5-line iPSCs using the Extracellular Analysis panel. Following four initial baseline measurements (file_1 to file_4), the concentration of quinidine was sequentially increased in the following sequence: 0.1 µM (file_5), 0.3 µM (file_6), 1 µM (file_7), 3 µM (file_8), 10 µM (file_9). Each data point in the provided figures corresponds to a value obtained from a single recording electrode. The horizontal and vertical bars denote the mean values and standard deviations (mean ± standard deviation), respectively.

We subsequently assessed the responses of SQT5-line CMs, subjected to other drugs listed in Table 2, while focusing on the drugs’ efficacy in prolonging the FPD. Both compounds, disopyramide (Figure S 2) and sotalol (Figure S 3), caused a dose-dependent prolongation of FPDs in CMs derived from SQT5 patients. On the other hand, flecainide (Figure S 4), ivabradine (Figure S 5), amiodarone (Figure S 6), and ranolazine (Figure S 7) did not or only minimally affect the FPD. These observations indicate that disopyramide and sotalol could be explored as alternative therapeutic options for treating arrhythmias in SQT5 patients, especially when quinidine proves ineffective or is unavailable.

Next, in the Intracellular Analysis panel of the CardioMEA Dashboard, we delved into the drug-induced modulations observed in the features of intracellular-like signals. Figure 9 illustrates the evolution of AP amplitude, depolarization time, APD50, and APD90 measured from iCell Cardiomyocytes, when the cells were exposed to nifedipine following three baseline measurements. Nifedipine is a Ca^2+^ channel blocker that shortens APDs [40]. As anticipated, nifedipine did not result in any substantial alteration of the AP amplitude or depolarization time. However, it significantly shortened APD_50_ and APD_90_, as evident from the visualization panel. A reduction in Ca^2+^ current renders the K^+^ current dominating during cardiac repolarization, leading to an abbreviated APD.

**Figure 9.**
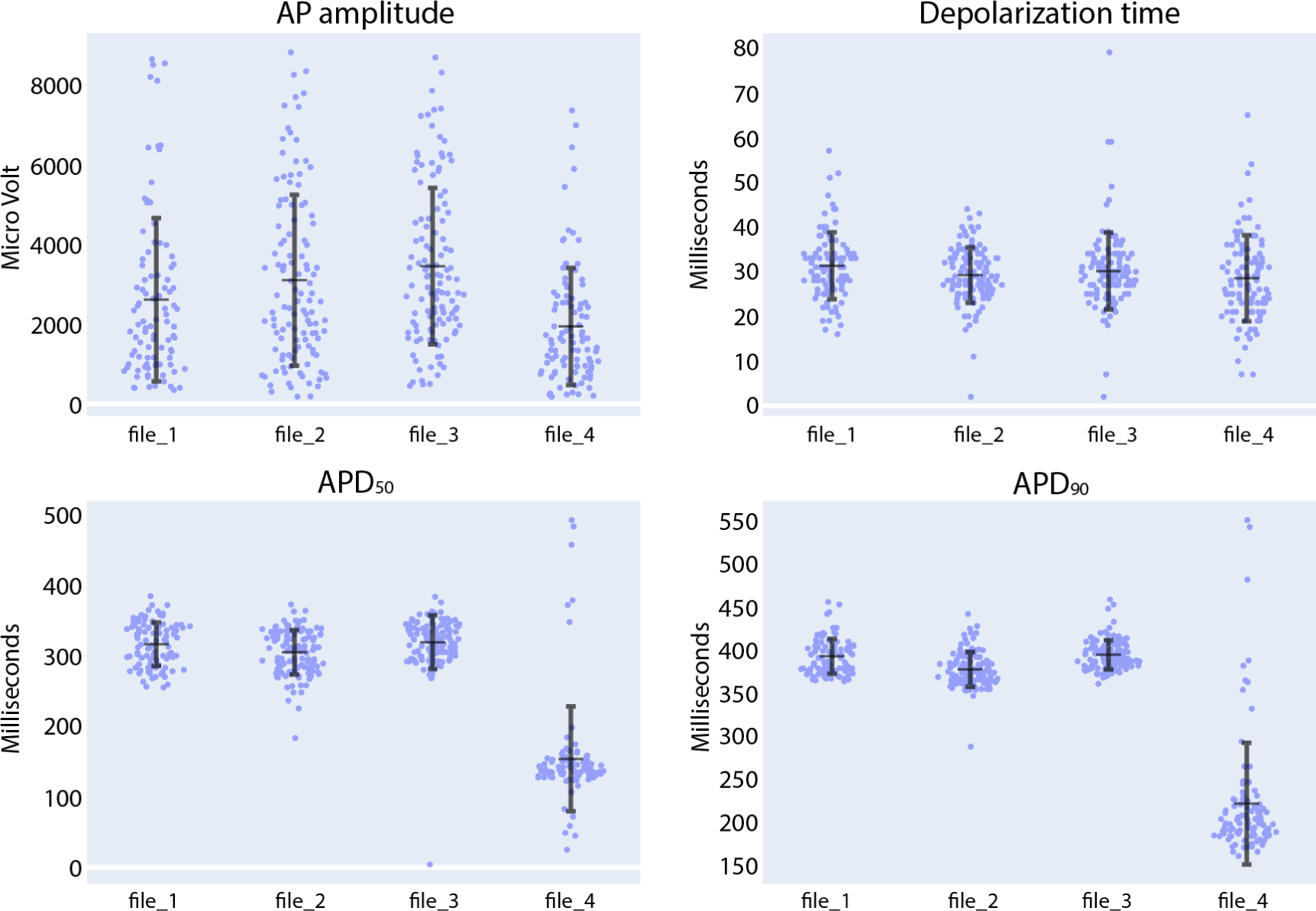
Compound analysis of signals of iCell Cardiomyocytes using the Intracellular Analysis panel. Following three initial baseline measurements (file_1 to file_3), nifedipine was added to the culture to reach a concentration of 100 nM (file_4). Each data point in the provided figures corresponds to a value obtained from a single recording electrode. The horizontal and vertical bars denote the mean value and standard deviation (mean ± standard deviation), respectively.

We further explored the effects of quinidine and sotalol on the AP features of iCell Cardiomyocytes. As shown in Figure S 8, exposure of CMs to 1 µM of quinidine increased depolarization time and APD_90_. The Na^+^ channel-blocking capability of quinidine and its effects on FPD prolongation have been previously documented[34, 35]. When subjected to 30 µM of sotalol (Figure S 9), there was negligible change in depolarization time [41], but we noted a notable rise in both APD_50_ and APD_90_. These observations are consistent with sotalol’s properties as a Class III, K^+^ channel blocker. As demonstrated by the analysis of the effects of seven clinically used drugs on extracellular signals and of three drugs on intracellular-like signals, CardioMEA holds promise as a versatile platform for data analysis and visualization in the field of MEA-based drug testing.

The second tab of the visualization and analysis panel, ‘Recording Info’, displays a table showing recording information and other features of the selected data files (Figure S 10). The third tab, exclusively available for the Extracellular Analysis panel and titled ‘Feature Analysis’, will be discussed in the subsequent section.

### Feature Analysis Using Automated Machine Learning

To identify key electrophysiological phenotypes and disease biomarkers that can be used to distinguish diseased cell lines from healthy controls, it is crucial to determine which features have a high predictive potential. To address this point, we incorporated automated machine learning (AutoML) and feature-importance analysis tools into CardioMEA, enabling users to run these complex analyses through the dashboard without the need to write any code.

In the Feature Analysis tab, users can select a subset of features from the data selected in the data panel with the assistance of a correlation heatmap, multicollinearity plot, and similarity cluster map (shown in Figure 10). Numerous features are extracted during the feature extraction process, and it is crucial to select a subset of these features before constructing classification models to discern diseased and healthy cell lines. This selection process is important, because some features may be highly correlated, exhibit high collinearity, or may be similar to each other, which may potentially interfere with the feature importance analysis. For example, permutation analysis [42], one of the essential techniques for investigating feature importance [43], could be affected, as collinear or similar features could compensate for permuting a feature. As a result, classification performance may not decline when one of the collinear features is permuted.

**Figure 10.**
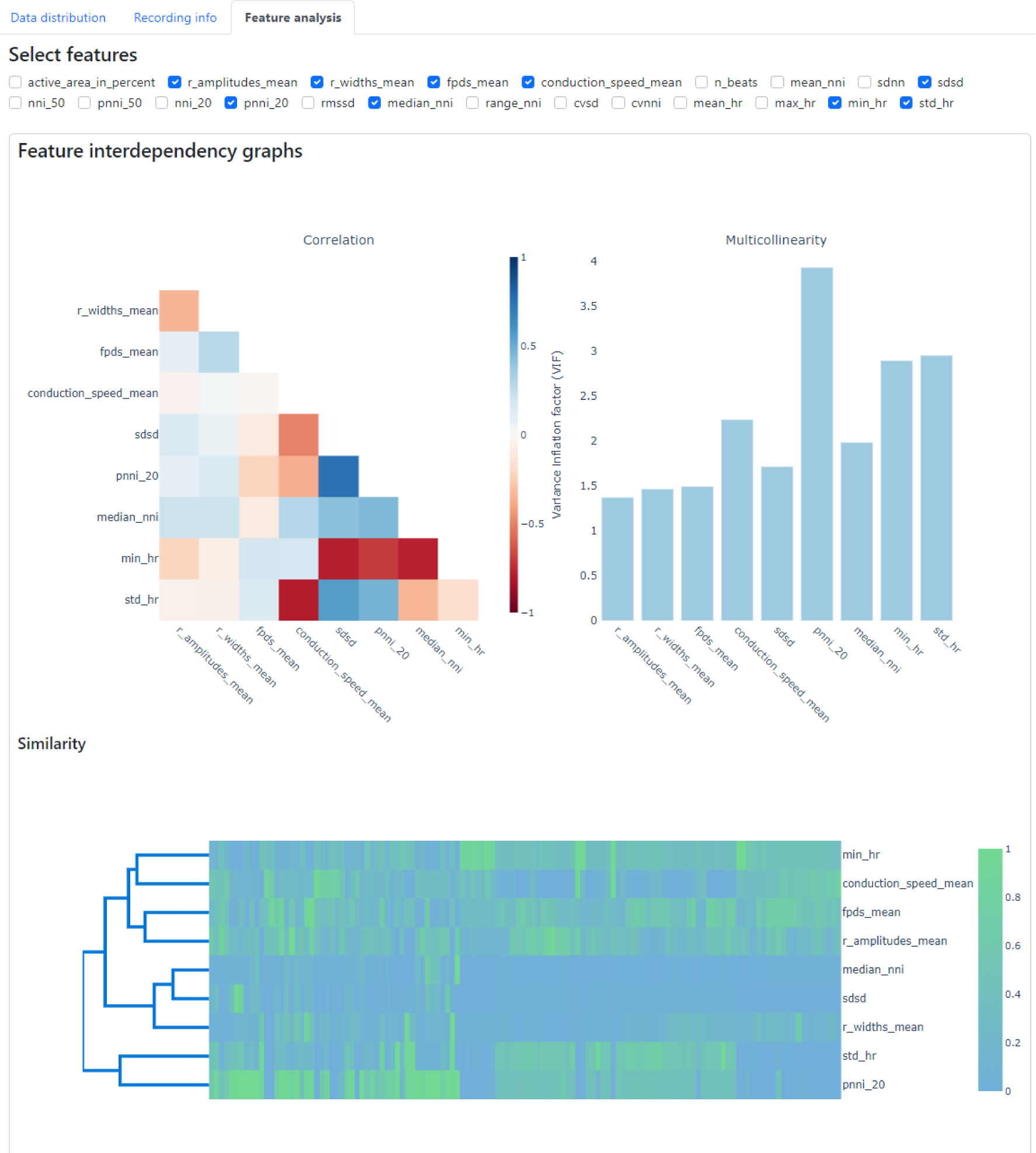
Feature analysis tab within the CardioMEA Dashboard, showing feature interdependency graphs. The correlation, multicollinearity, and similarity between selected features were computed and displayed.

To demonstrate the usage of Feature Analysis, we selected recording data from all cultures without drug administration in the data panel to investigate which features are crucial in distinguishing SQT5-line and SQT5corr-line CMs. In total, 93 cultures of SQT5- line CMs and 33 cultures of SQT5corr-line CMs were included for feature importance analysis.

Using the Feature interdependency graphs in the panel, we eliminated features from the list, which were highly correlated, clustered as similar, or had pronounced multicollinearities. Figure 10 presents a screenshot of the dashboard showing the selected 9 features after eliminating redundant features using a correlation heatmap, multicollinearity plot, and similarity cluster map. This screenshot demonstrates CardioMEA’s unique advantage of assisting the user in selecting distinctive features and eliminating redundant ones in a user-friendly and interactive manner.

After the selection of features, the ensuing step involves investigating their importance. Within CardioMEA, we have integrated an AutoML functionality to identify the optimal model for classification purposes. This functionality has been realized by harnessing the AUTO-SKLEARN toolkit [44], an open-source library that facilitates an efficient and automated approach to machine learning model selection and hyperparameter tuning. The user is empowered to determine how missing data is handled and to set the test data size, the number of cross-validation folds, the time limit per fold, and the number of permutation repeats (Figure 11). By simply clicking ‘Run AutoML’, the AutoML algorithm is triggered.

**Figure 11.**
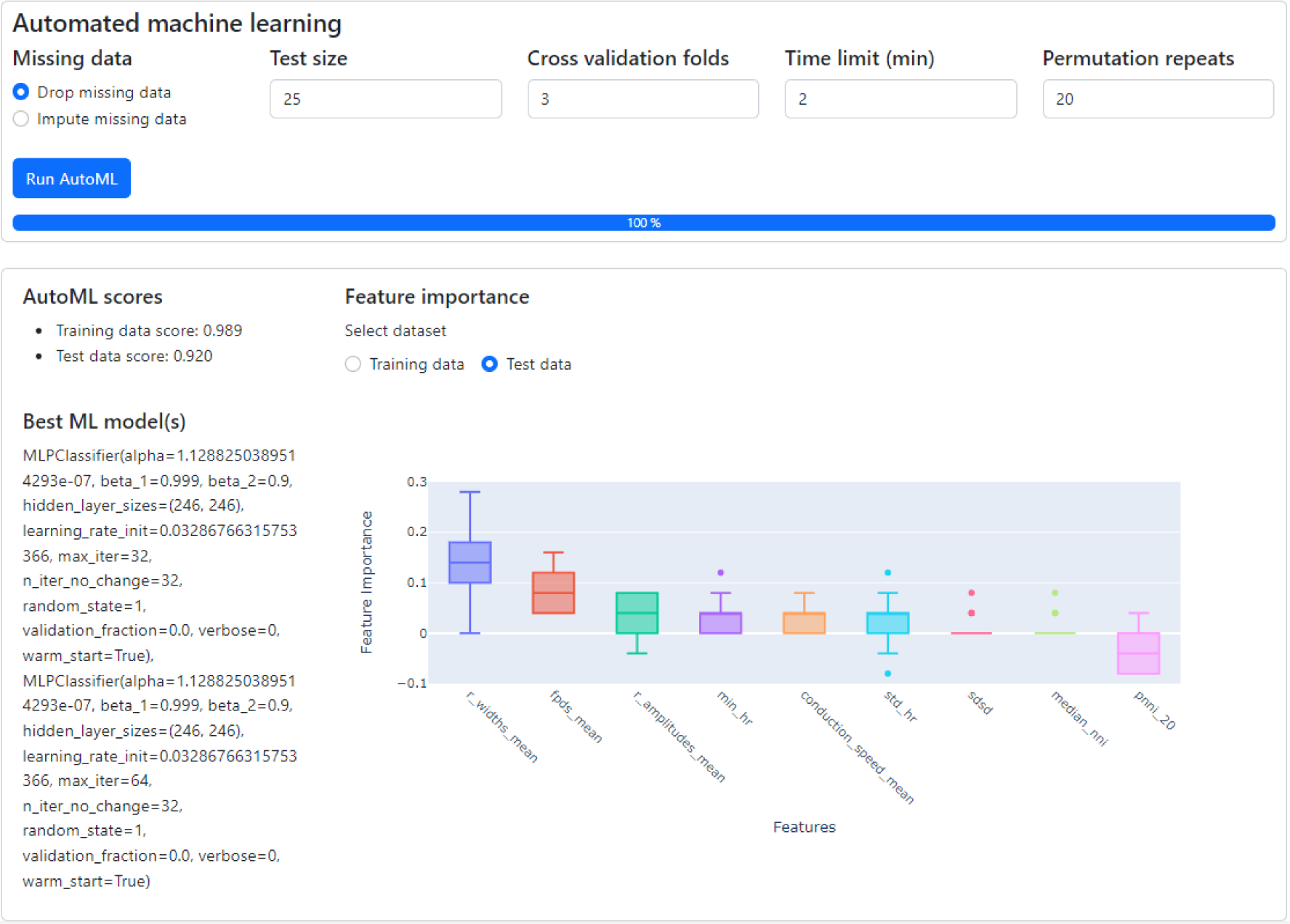
The classification of CMs derived from SQT5-line and SQT5corr-line iPSCs, followed by an analysis of feature importance. With just a few mouse clicks, users are empowered to perform automated machine learning to build the optimal classification model. The performance and details of the optimized model are displayed in the lower left section, along with the outcomes of a feature importance analysis (lower right section). Given the respective ML model, this analysis shows the predictive power of selected features for distinguishing between SQT5-line and SQT5corr-line CMs.

As shown in the lower left section of Figure 11, the AutoML process yielded the result that the multi-layer perceptron (MLP) classifier delivered the best performance for the provided dataset, applying a 3-fold, stratified cross-validation (sCV) method. This model achieved a classification accuracy of 98.9 % for the training dataset and 92.0 % for the test dataset. We compared this result with the outcomes of a baseline model, which classified the data based on the most frequent labels. Upon conducting this procedure 10 times with a 5-fold sCV approach, the baseline model yielded an accuracy of 71.8 ± 1.8 % (mean ± standard deviation). Next, employing the MLP classifier with its default parameters in the SKLEARN library [45] yielded 80.1 ± 5.3 % accuracy. The comparison between the baseline model and the default MLP classifier with the model constructed by AutoML shows that AutoML identifies and fine-tunes an optimal model, thereby enhancing performance.

Subsequently, CardioMEA computes feature importance values using the optimized model. This process involves permutating the values of each feature individually and gauging the subsequent decline in accuracy over a predefined set of iterations [42]. As depicted in Figure 11 in the lower right section, R-wave spike width and FPD were identified as the top two significant features. This finding suggests that these features play a pivotal role in the classification of SQT5-line and SQT5corr-line cells when utilizing the AutoML-trained model. It is crucial, however, to emphasize that permutation importance indicates the significance of a specific feature for a particular model, which is the MLP classifier in this case, rather than representing its innate predictive power. As demonstrated in the context of Figure 11, users can efficiently construct optimal machine learning models using the dashboard, bypassing the need to manually code the algorithms, which significantly simplifies the process of identifying critical features for distinguishing between diseased cell lines and healthy controls.

The aforementioned analysis can be conducted on the dashboard with just a few mouse clicks. Users do not need to delve into the intricate details of constructing machine learning models or analyzing feature importance when exploring their HD-MEA data collected from CMs. This feature demonstrates that the CardioMEA Dashboard can serve as a user-friendly, “no-code” platform for advanced data analysis and visualization.

We further investigated two features, which emerged as the top two determinants in distinguishing between diseased and healthy cell lines in our feature importance analysis, the R-wave spike width and FPD. Given that the SQT5corr-line was generated by correcting the CACNB2 gene of SQT5, the feature analysis suggests that the mutation within the CACNB2 gene of the SQT5 cells may alter the R-wave spike width and FPD values. A previous study [28] revealed that CMs, differentiated from the same SQT5-line iPSCs, exhibited decreased Na^+^ and L-type Ca^2+^ channel currents. Figure 12 shows a statistical comparison of these feature values - R-wave spike width and FPD - between SQT5-line and SQT5corr-line CM cultures.

**Figure 12.**
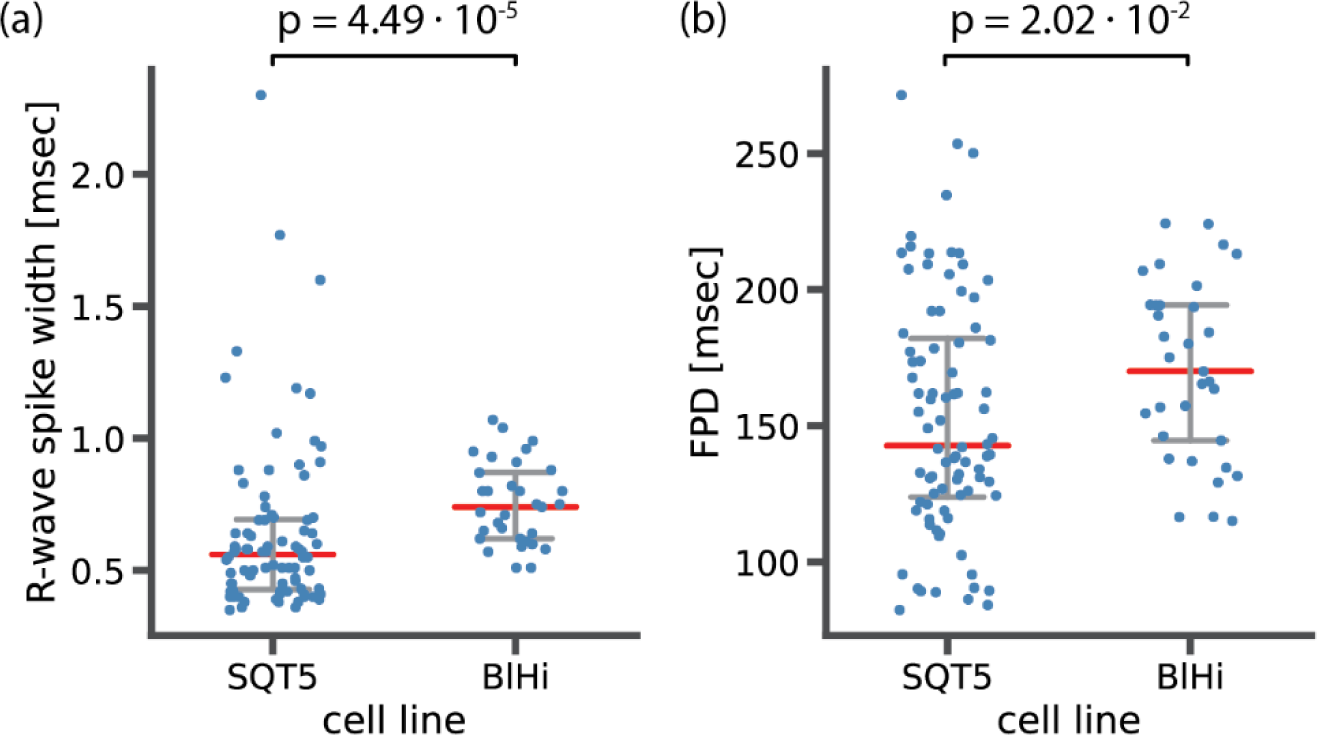
Statistical analysis to compare (a) R-wave spike width and (b) field potential duration (FPD) between diseased (SQT5) and healthy control (SQT5corr) cell lines. Each data point (blue color) represents one CM culture. Red bars indicate median values and error bars (grey color) indicate interquartile ranges. The p values were estimated using a two-sided Mann-Whitney U test.

As shown in Figure 12a, the observed median value of the R-wave spike width was lower in SQT5 cultures than in SQT5corr cultures. We had anticipated that the R-wave spike width would be larger in SQT5 than in SQT5corr cultures, given that reduced I_Na_ should result in a slower depolarization. However, the range of measured data points obtained from SQT5 cultures is significantly larger than that of SQT5corr cultures. This large range renders it difficult to draw solid conclusions from statistical analysis. Although the statistical analysis did not provide a conclusive insight into the relationship of R-wave spike width values between the SQT5 and SQT5corr cultures, the feature importance analysis, carried out via the CardioMEA Dashboard, as shown in Figure 11, identified the R-wave spike width as the most predictive feature. Unlike traditional statistical comparisons, the machine learning algorithm - an MLP classifier in this case - utilizes all provided features collectively to distinguish between SQT5 and SQT5corr cultures. The results of the feature analysis evidence the potential of CardioMEA’s AutoML-driven module. This module could improve the identification of novel disease phenotypes, an area where standalone statistical analysis may fall short.

As shown in Figure 12b, FPD values were lower in the SQT5 cultures compared to the SQT5corr cultures. Given the established understanding that mutations in the CACNB2 gene in the SQT5 cell line induce a loss of function in L-type Ca^2+^ channels [28], this decrease of FPD in SQT5 cultures was anticipated. Therefore, the FPD was identified as the second most influential feature in identifying SQT5 cultures, yielding a statistically significant difference between the SQT5 and SQT5corr cultures.

## Conclusions

The study of therapeutic efficacy and potential cardiotoxicity of drugs *in vitro* is a crucial step of preclinical research. HD-MEAs, with their high SNR and spatiotemporal resolution, are instrumental in characterizing the electrophysiology of iPSC-derived CMs. However, until now, there has been a lack of open-source data analysis platforms that can cope with the large data volumes generated by HD-MEA technology.

In this study, we presented CardioMEA, a comprehensive data analysis platform with the ability to process and analyze large volumes of CM data generated by HD-MEAs. CardioMEA offers a complete analysis workflow, from data extraction to exploratory analysis. The platform’s first component provides scalable data processing for feature extraction across multiple data files, while the second component consists of a user-friendly, interactive web-based dashboard for advanced data visualization and analysis. Using CardioMEA, we examined the efficacy of seven clinically used drugs on CMs derived from an SQT5 patient and the cardiotoxicity of three drugs on healthy-donor-derived CMs.

The feature analysis tool within the CardioMEA Dashboard assists users in feature selection and helps to reduce redundancy and multicollinearity in the data. Furthermore, AutoML and feature importance analysis provide insights into the predictive power of features for discerning electrophysiological signatures of diseased and healthy CMs. CardioMEA allows users to perform complex analyses on their HD-MEA data without extensive coding knowledge.

Another unique feature of CardioMEA is its ability to process intracellular signals captured by HD-MEAs, which renders CardioMEA the first of its kind among open-source platforms. The possibility to process both extracellular and intracellular signals renders the platform amenable to a wide range of researchers’ potential needs.

Overall, CardioMEA is a comprehensive and user-friendly platform that significantly advances the analysis of CM signals originating from HD-MEAs. The combination of parallel data processing, interactive dashboard, and feature analysis tools enables users to explore and analyze their data efficiently. The platform is expected to advance studies of cardiac diseases and drug testing and to democratize associated high-level data analysis.

## Author Contributions

Conceptualization, J.L., I.E., A.M.S., F.D., V.E., A. Hierlemann and H.U.; experiment and data collection, J.L., E.D.; preparation and provision of cell lines, I.E., A. Hohn and L.C.; data analysis, J.L. and E.D.; funding acquisition, F.D. and A. Hierlemann; writing−original draft, J.L.; writing−review and editing, E.D., I.E., A. Hohn, A.M.S., F.D., L.C., A. Hierlemann and H.U.; All authors have read and agreed to the published version of the manuscript.

## Funding

This research was supported by the Personalized Health and Related Technologies initiative of the ETH Domain (Project No. PHRT-iDoc 2018-329) and the Swiss National Science Foundation under Grant 205320_188910/1.

## Ethics Statement

The generation of iPSCs was approved by the Ethics Committee of the University Medical Center Göttingen (approval number: 10/9/15) and carried out in accordance with the approved guidelines. Written informed consent was obtained from all participants or their legal representatives prior to the participation in the study.

## Acknowledgements

The authors are grateful to Dr. Taehoon Kim for providing scientific inputs to the data analysis. The authors are particularly thankful to Dr. Xiaohan Xue and Dr. Marc Emmenegger for their feedback on this Manuscript. They also wish to acknowledge the High Performance Computing group for operating and maintaining the Euler cluster at ETH Zurich, which was used for parallelized processing of the massive volume of data in this study. The authors used ChatGPT version 4 from OpenAI for English language editing.

## Conflict of Interest

A.M.S. received educational grants through his institution from Abbott, Bayer Healthcare, Biosense Webster, Biotronik, Boston Scientific, BMS/Pfizer, and Medtronic; and speaker /advisory board /consulting fees from Bayer Healthcare, Biotronik, Medtronic, Novartis, Pfizer, Stride Bio Inc., and Zoll Medical. The other authors declare no conflict of interest.

