## supplementary information for "CardioMEA: Comprehensive Data Analysis Platform for Studying Cardiac Diseases and Drug Responses"

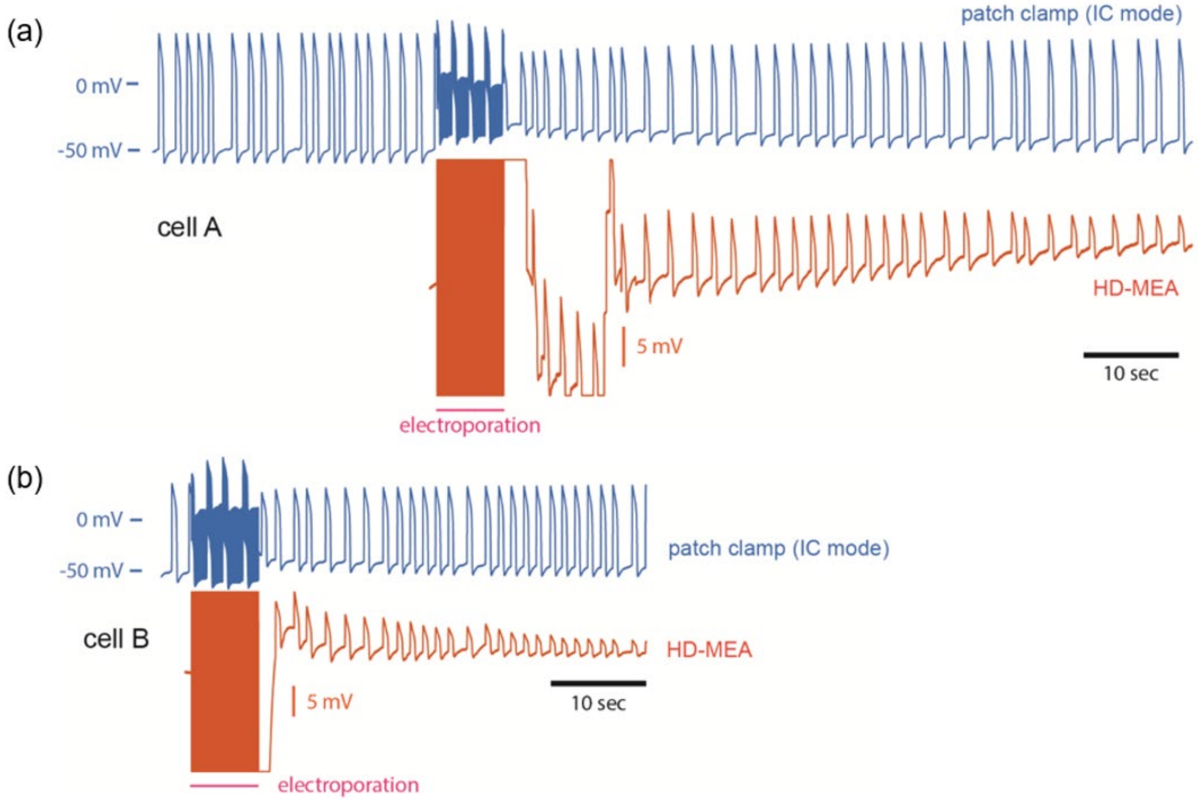


Figure S 1. Intracellular recordings of (a) cell A and (b) cell B by whole-cell patch clamp, in the current-clamp mode (blue) and intracellular-like recordings obtained by the HD-MEA upon electroporation (red), obtained simultaneously. Reprinted with permission from ACS Sensors 2022, 7, 10, 3181–3191. Copyright 2022 American Chemical Society.

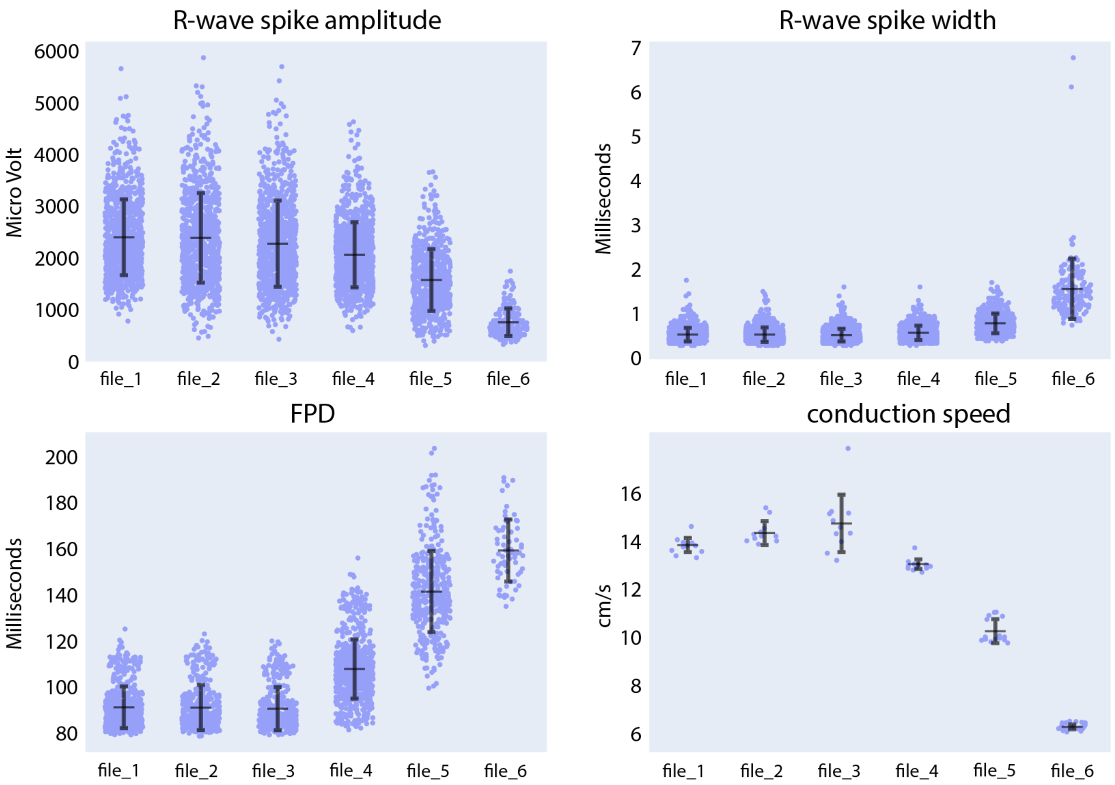


Figure S 2. Compound analysis of signals of CMs derived from SQT5-line iPSCs using the Extracellular Analysis panel. Following four initial baseline measurements (file_1 to file_3), the concentration of disopyramide was sequentially increased in the following sequence: 3 µM (file_4), 13 µM (file_5), 43 µM (file_6). Each data point in the provided figures corresponds to a value obtained from a single recording electrode. The horizontal and vertical bars denote the mean values and standard deviations (mean ± standard deviation), respectively.

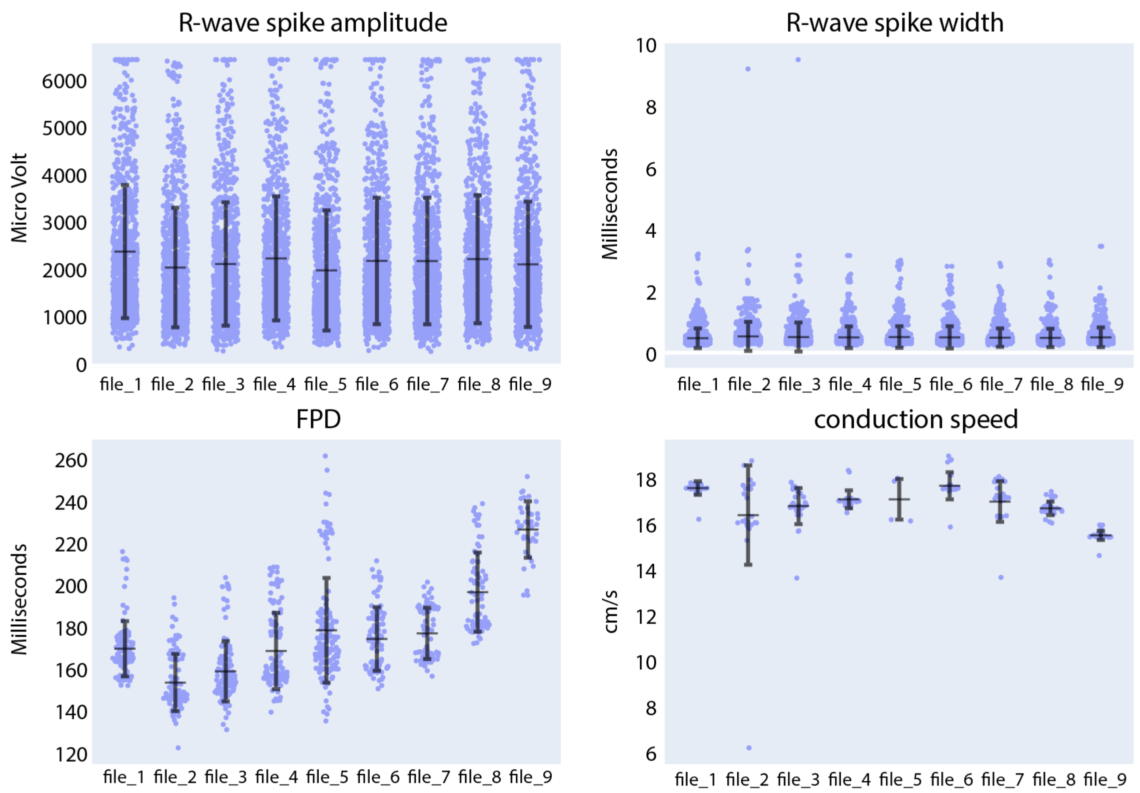


Figure S 3. Compound analysis of signals of CMs derived from SQT5-line iPSCs using the Extracellular Analysis panel. Following four initial baseline measurements (file_1 to file_4), the concentration of sotalol was sequentially increased in the following sequence: 3 µM (file_5), 10 µM (file_6), 30 µM (file_7), 100 µM (file_8), 300 µM (file_9). Each data point in the provided figures corresponds to a value obtained from a single recording electrode. The horizontal and vertical bars denote the mean values and standard deviations (mean ± standard deviation), respectively.

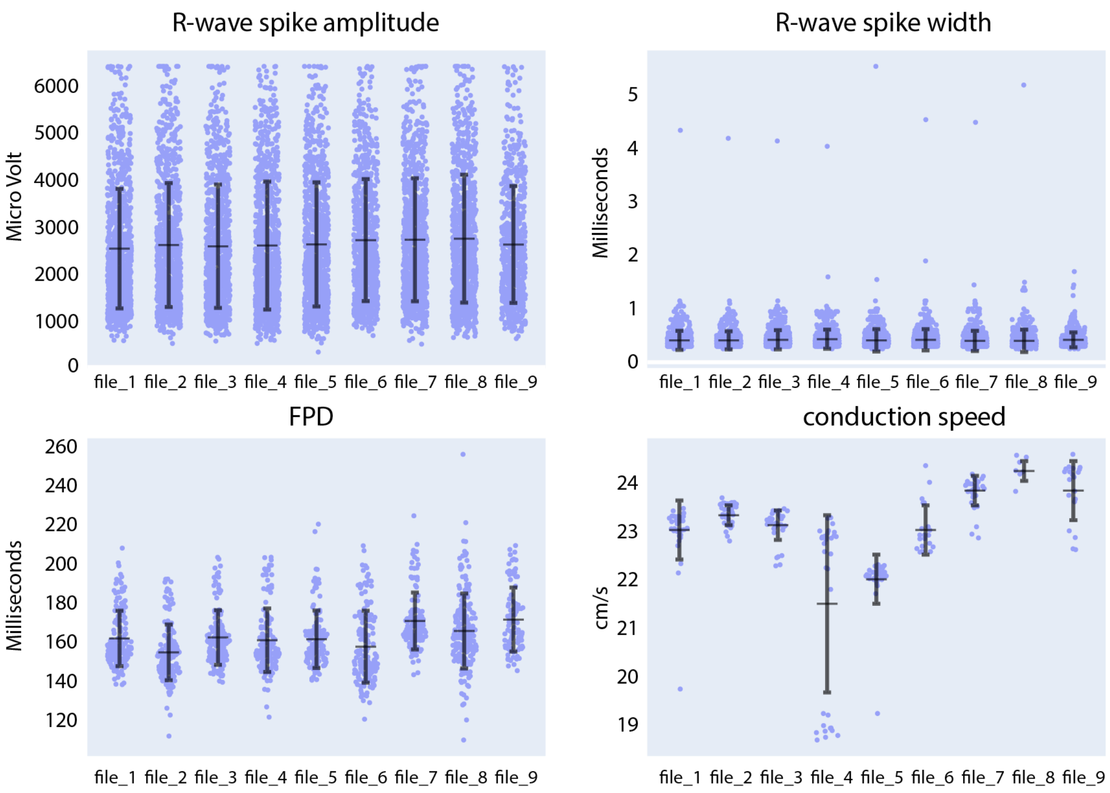


Figure S 4. Compound analysis of signals of CMs derived from SQT5-line iPSCs using the Extracellular Analysis panel. Following four initial baseline measurements (file_1 to file_4), the concentration of flecainide was sequentially increased in the following sequence: 0.03 µM (file_5), 0.1 µM (file_6), 0.3 µM (file_7), 1 µM (file_8), 3 µM (file_9). Each data point in the provided figures corresponds to a value obtained from a single recording electrode. The horizontal and vertical bars denote the mean values and standard deviations (mean ± standard deviation), respectively.

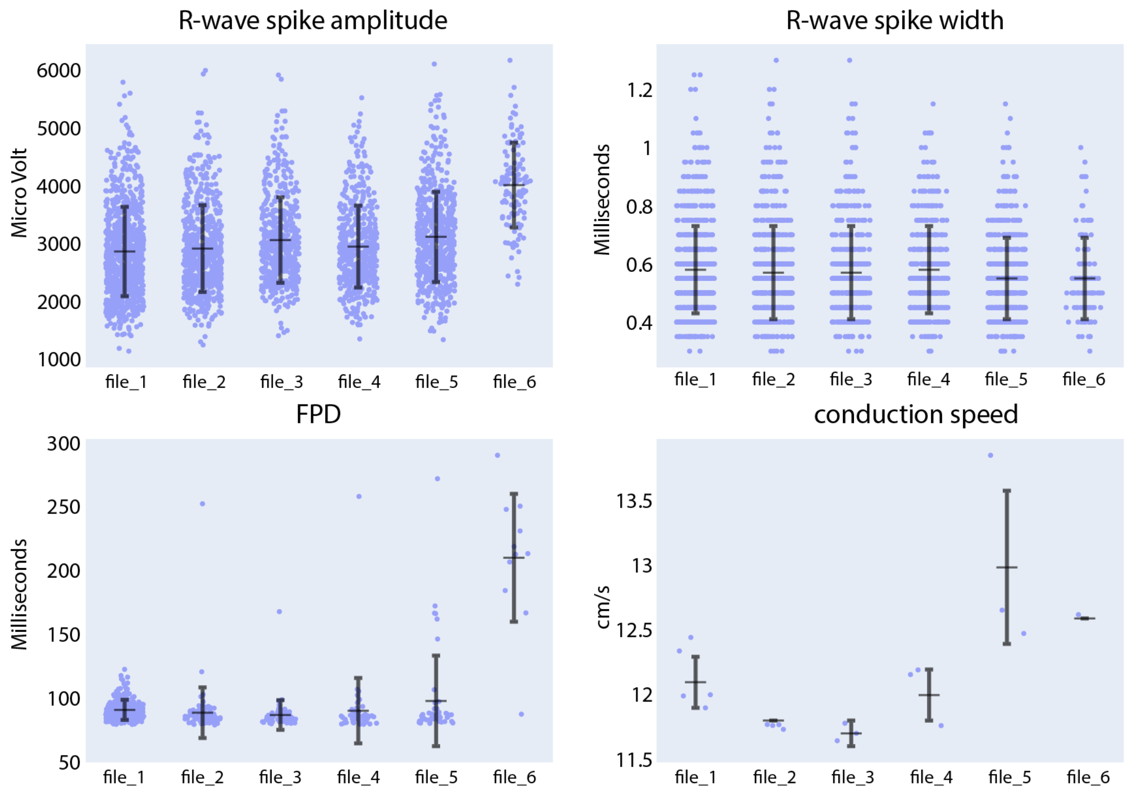


Figure S 5. Compound analysis of signals of CMs derived from SQT5-line iPSCs using the Extracellular Analysis panel. Following four initial baseline measurements (file_1 to file_3), the concentration of ivabradine was sequentially increased in the following sequence: 0.3 µM (file_4), 1 µM (file_5), 3 µM (file_6). Each data point in the provided figures corresponds to a value obtained from a single recording electrode. The horizontal and vertical bars denote the mean values and standard deviations (mean ± standard deviation), respectively.

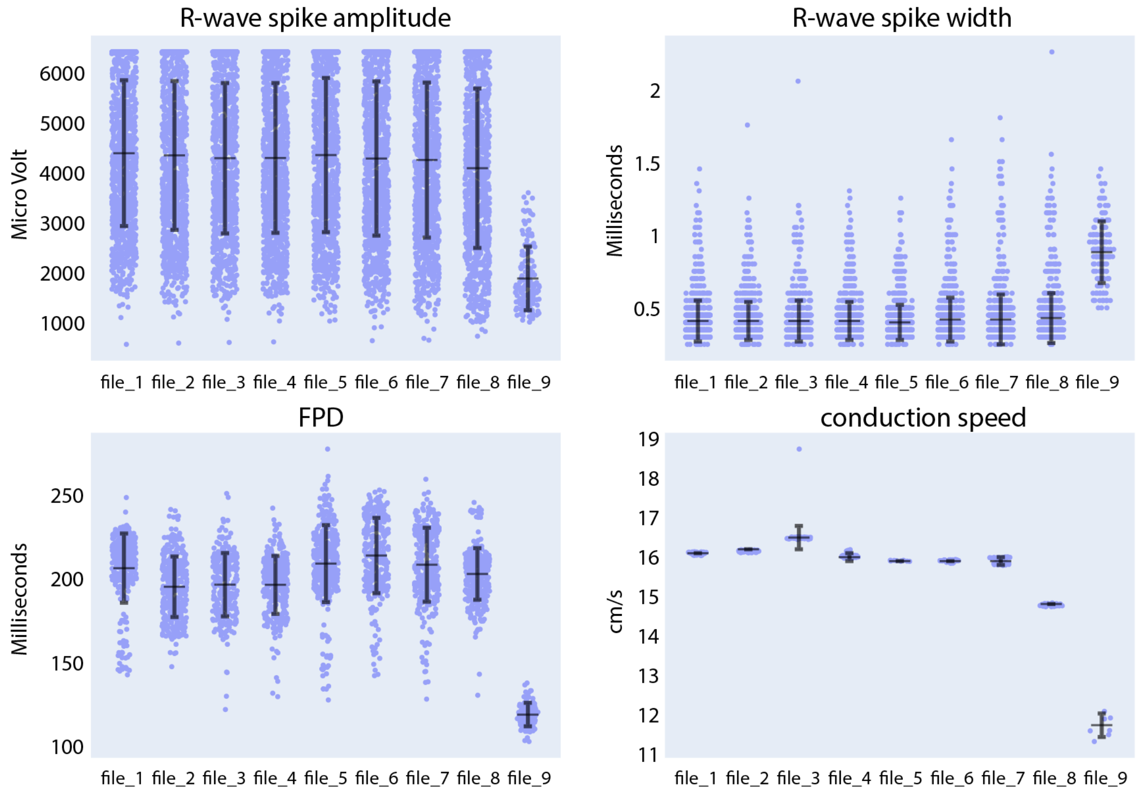


Figure S 6. Compound analysis of signals of CMs derived from SQT5-line iPSCs using the Extracellular Analysis panel. Following four initial baseline measurements (file_1 to file_4), the concentration of amiodarone was sequentially increased in the following sequence: 0.03 µM (file_5), 0.1 µM (file_6), 0.3 µM (file_7), 1 µM (file_8), 3 µM (file_9). Each data point in the provided figures corresponds to a value obtained from a single recording electrode. The horizontal and vertical bars denote the mean values and standard deviations (mean ± standard deviation), respectively.

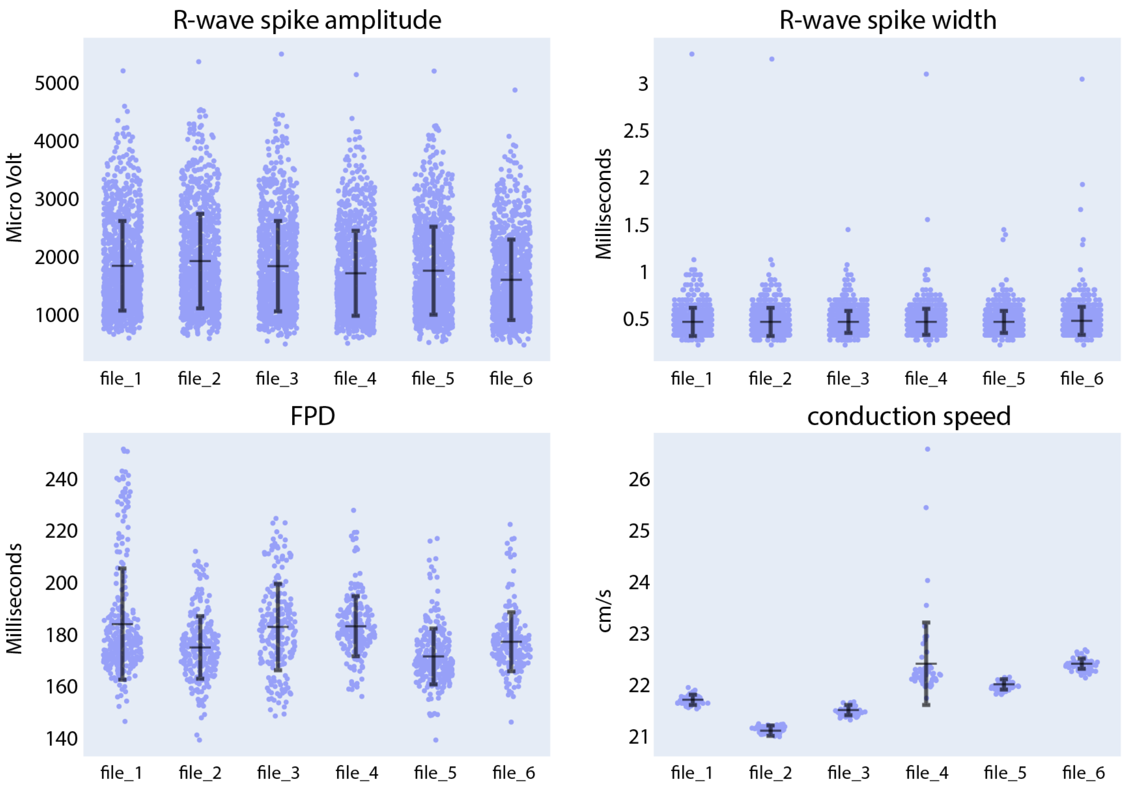


Figure S 7. Compound analysis of signals of CMs derived from SQT5-line iPSCs using the Extracellular Analysis panel. Following four initial baseline measurements (file_1 to file_3), the concentration of ranolazine was sequentially increased in the following sequence: 0.3 µM (file_4), 1 µM (file_5), 3 µM (file_6). Each data point in the provided figures corresponds to a value obtained from a single recording electrode. The horizontal and vertical bars denote the mean values and standard deviations (mean ± standard deviation), respectively.

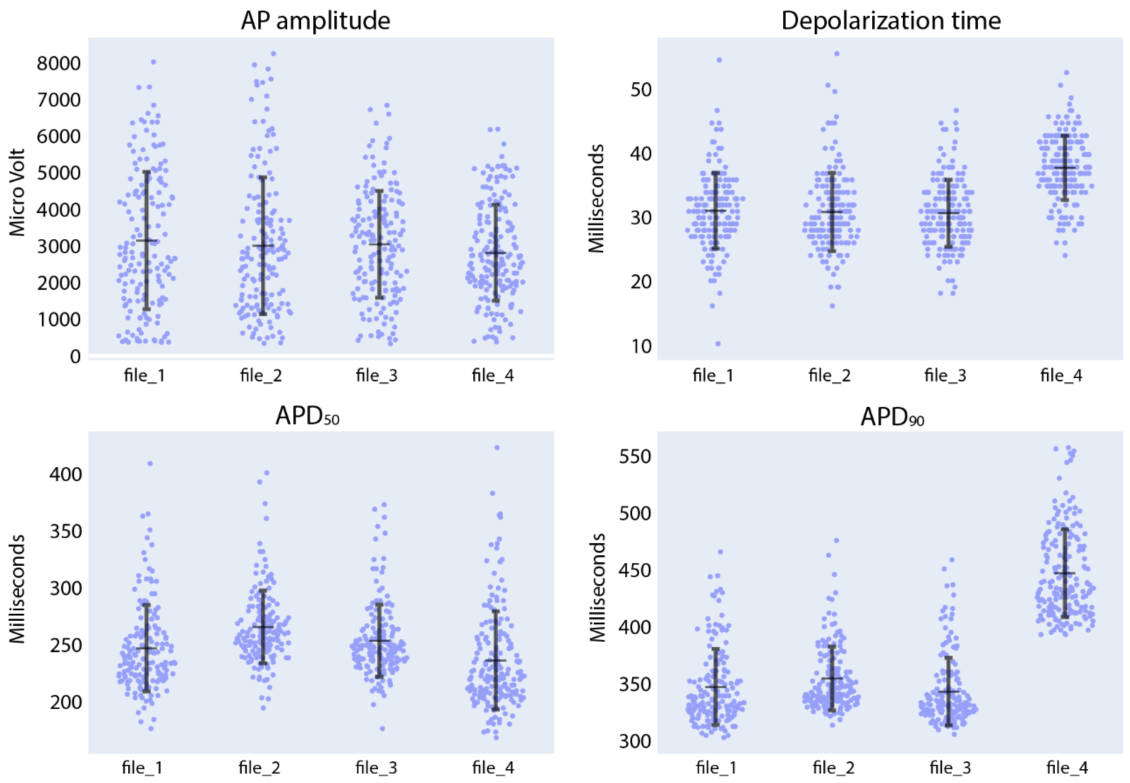


Figure S 8. Compound analysis of signals of iCell Cardiomyocytes using the Intracellular Analysis panel. Following three initial baseline measurements (file_1 to file_3), quinidine was added to the culture to reach a concentration of 1 µM (file_4). Each data point in the provided figures corresponds to a value obtained from a single recording electrode. The horizontal and vertical bars denote the mean value and standard deviation (mean ± standard deviation), respectively.

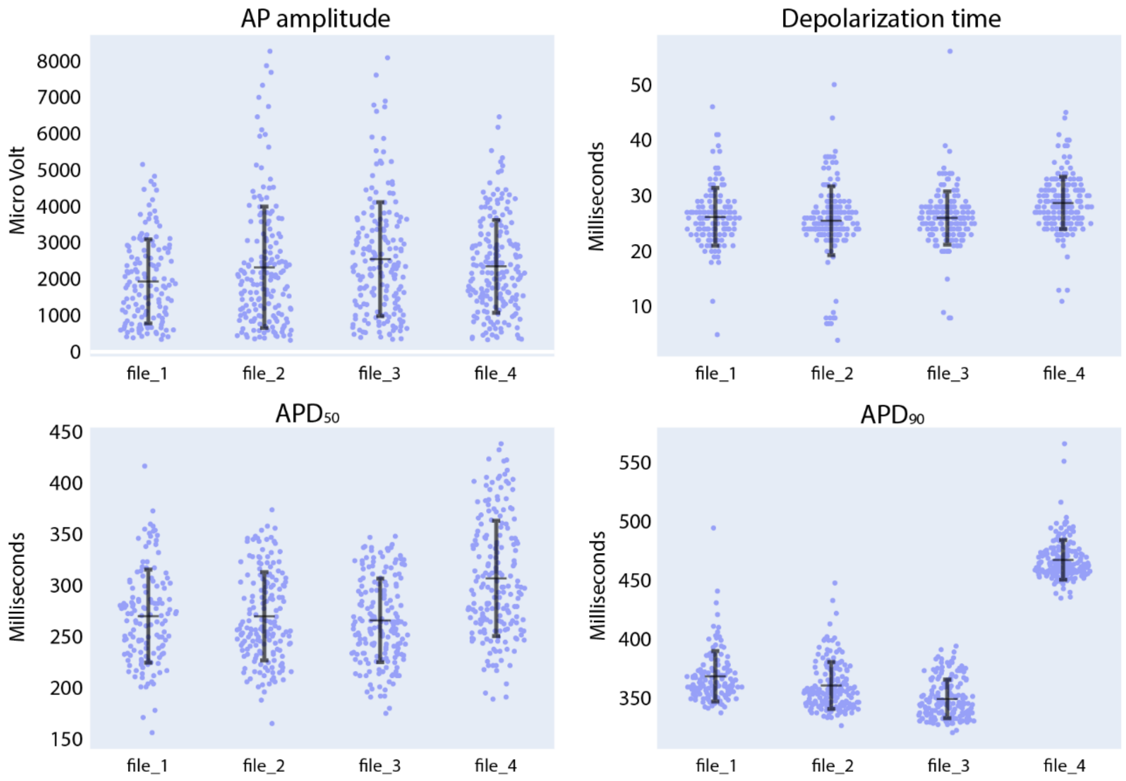


Figure S 9. Compound analysis of signals of iCell Cardiomyocytes using the Intracellular Analysis panel. Following three initial baseline measurements (file_1 to file_3), sotalol was added to the culture to reach a concentration of 30 µM (file_4). Each data point in the provided figures corresponds to a value obtained from a single recording electrode. The horizontal and vertical bars denote the mean value and standard deviation (mean ± standard deviation), respectively.

Data distribution | **Recording info** | Feature analysis

| index | file_1 | file_2 | file_3 | file_4 | file_5 | file_6 |
|---|---|---|---|---|---|---|
| gain | 512 | 512 | 512 | 512 | 512 | 512 |
| n_electrodes_sync | 957 | 875 | 919 | 922 | 594 | 165 |
| active_area_in_percent | 93.8 | 85.8 | 90.1 | 90.4 | 58.2 | 16.2 |
| rec_duration | 59.1 | 60.1 | 60.1 | 120.1 | 120.1 | 119.4 |
| rec_proc_duration | 59.1 | 60 | 60 | 60 | 60 | 60 |
| n_beats | 11 | 11 | 11 | 12 | 15 | 19 |
| mean_nni | 5107.1 | 5227.9 | 5702.4 | 5091.4 | 3937.9 | 3098.1 |
| sdnn | 180.4 | 144.1 | 788.2 | 207.5 | 917.4 | 77.9 |
| sdsd | 180.7 | 132.5 | 1088.6 | 326.5 | 1607.1 | 121.4 |
| nni_50 | 7 | 6 | 9 | 10 | 13 | 12 |
| pnni_50 | 77.8 | 66.7 | 100 | 100 | 100 | 70.6 |
| nni_20 | 7 | 8 | 9 | 10 | 13 | 13 |
| pnni_20 | 77.8 | 88.9 | 100 | 100 | 100 | 76.5 |
| rmssd | 184.3 | 137.4 | 1091.2 | 326.7 | 1607.1 | 121.4 |
| median_nni | 5079.1 | 5251 | 5533.3 | 4993 | 3438.9 | 3079.2 |
| range_nni | 514.9 | 529.1 | 2922 | 505.9 | 2172.3 | 331.6 |
| cvsd | 0 | 0 | 0.2 | 0.1 | 0.4 | 0 |
| cvnni | 0 | 0 | 0.1 | 0 | 0.2 | 0 |
| mean_hr | 11.8 | 11.5 | 10.7 | 11.8 | 15.9 | 19.4 |
| max_hr | 12.3 | 12.2 | 12.4 | 12.3 | 19.1 | 20.4 |
| min_hr | 11.1 | 11 | 7.7 | 11.2 | 11.3 | 18.3 |
| std_hr | 0.4 | 0.3 | 1.1 | 0.5 | 3.2 | 0.5 |

Figure S 10. Detailed information on each recording file can be found under the "Recording info" tab in the CardioMEA Dashboard. The image shows data generated during disopyramide administration to CMs derived from SQT5-line iPSCs. Following four initial baseline measurements (file_1 to file_3), the concentration of disopyramide was sequentially increased in the following sequence: 3 µM (file_4), 13 µM (file_5), 43 µM (file_6).
